# Defense-related callose synthase *PMR4* promotes root hair callose deposition and adaptation to phosphate deficiency in *Arabidopsis thaliana*

**DOI:** 10.1101/2023.07.05.547890

**Authors:** Kentaro Okada, Koei Yachi, Tan Anh Nhi Nguyen, Satomi Kanno, Shigetaka Yasuda, Haruna Tadai, Chika Tateda, Tae-Hong Lee, Uyen Nguyen, Kanako Inoue, Natsuki Tsuchida, Taiga Ishihara, Shunsuke Miyashima, Kei Hiruma, Kyoko Miwa, Takaki Maekawa, Michitaka Notaguchi, Yusuke Saijo

## Abstract

Plants acquire phosphorus (P) primarily as inorganic phosphate (Pi) from the soil. Under Pi deficiency, plants induce an array of physiological and morphological responses, termed phosphate starvation response (PSR), thereby increasing Pi acquisition and use efficiency. However, the mechanisms by which plants adapt to Pi deficiency remain to be elucidated. Here, we report that deposition of a β-1,3-glucan polymer called callose is induced in *Arabidopsis thaliana* root hairs under Pi deficiency, in a manner independent of PSR-regulating *PHR1/PHL1* transcription factors and *LPR1/LPR2* ferroxidases. Genetic studies revealed *PMR4* (*GSL5*) callose synthase being required for the callose deposition in Pi-depleted root hairs. Loss of *PMR4* also reduces Pi acquisition in shoots and plant growth under low Pi conditions. The defects are not recovered by simultaneous disruption of *SID2*, mediating defense-associated salicylic acid (SA) biosynthesis, excluding SA defense activation from the cause of the observed *pmr4* phenotypes. Grafting experiments and characterization of plants expressing *PMR4* specifically in root hair cells suggest that a PMR4 pool in the cell type contributes to shoot growth under Pi deficiency. Our findings thus suggest an important role for *PMR4* in plant adaptation to Pi deficiency.

**Significance statement:** We reveal that PMR4 callose synthase mediates callose deposition in root hairs under phosphate (Pi) deficiency, without requiring Pi starvation response regulators *PHR1/PHL1* or *LPR1/LPR2*. The loss of the callose deposition is accompanied by decreases in Pi acquisition and plant growth in *pmr4*. Root hair cell-specific *PMR4* expression restores callose deposition in root hairs and shoot growth under Pi deficiency, indicating a critical role for root hair callose in plant adaptation to Pi deficiency.

## Introduction

As an essential macronutrient, plants acquire and distribute P from the rhizosphere to the whole tissues, primarily in the form of inorganic phosphate (Pi). To maintain Pi homeostasis under Pi deficiency, plants induce an array of adaptive responses, *e.g.* increasing Pi uptake, remobilization and utilization, termed Pi starvation response (PSR), and involve beneficial microbes in mutualistic associations (Péret et al., 2011; Chien et al., 2018; Isidra-Arellano et al., 2021). PSR is accompanied by extensive transcriptional reprogramming, mainly through the transcription factor *PHR1* and related *PHL1*, which leads to increases in Pi transporter expression, root hair formation and anthocyanin accumulation (Rubio et al., 2001, Bustos et al., 2010). *PHR1/PHL1* mediate systemic coordination of these responses according to the internal Pi availability. In addition, Pi depletion leads to remodeling root system architecture, a decrease in primary root growth while increasing lateral root formation and growth, through the ferroxidases *LPR1* and *LPR2* (Svistoonoff et al., 2007; Péret et al., 2014; Balzergue et al., 2017). *LPR1/LPR2* are required for arresting primary root growth when the external Pi levels are reduced surrounding the root tips (Svistoonoff et al., 2007).

Recent studies illuminated an intimate relationship between PSR and biotic interactions, but the molecular basis for their coordination remains poorly understood (Chan et al., 2021).

Callose deposition on the plasma membrane through callose synthases *Glucan Synthase Likes* (*GSLs*) reinforces barrier functions of the cell wall and restricts plasmodesmatal permeability, in plant development and biotic/abiotic stress responses (Jacobs et al., 2003; Nishimura et al., 2003; Dong et al., 2005; Enns et al., 2005; Nishikawa et al., 2005; Chowdhury et al., 2014; Ellinger & Voigt, 2014; Sager & Lee, 2014; Wu et al., 2018). In *Arabidopsis thaliana* (hereafter Arabidopsis), callose deposition is induced in the root phloem in response to excess iron, thereby reducing plasmodesmatal permeability (O’Lexy et al., 2018). Of 12 Arabidopsis *GSLs*, *GSL2* (*CALS5*) and *GSL5* (*PMR4, CALS12*) respectively contribute to basal accumulation and iron-induced deposition of callose in the root phloem (O’Lexy et al., 2018). Callose also accumulates in the stem cell niche (SCN) surrounding the quiescent center cells, thereby inhibiting cell-to-cell movement of the transcription factor SHORT ROOT (SHR) during PSR. This serves to suppress cell division and elongation in the root apical meristem and elongation zone, respectively, through the *LPR1/LPR2* pathway (Müller et al., 2015; Mora-Macías et al., 2017). *GSL5* (*PMR4*) also mediates callose deposition in response to pathogen challenge and microbe/damage-associated molecular patterns (MAMPs/DAMPs), and promotes fungal penetration resistance (Wang et al., 2021).

However, the loss of *GSL5* function results in strong resistance to powdery mildew fungi through tightened SA-based defenses, demonstrated by *gsl5* mutants revealed as *powdery mildew resistance4* (*pmr4*) (Vogel & Somerville, 2000; Nishimura et al., 2003).

This resistance is thought to represent a backup defense activated when *GSL5* (*PMR4*)-mediated callose deposition, a key output of MAMP/DAMP-triggered defense signaling, is disrupted.

Although the significance of *PMR4* has been genetically shown, our knowledge is very limited for the mechanisms that involve and regulate PRM4 in callose deposition during defense responses (Wang et al., 2021). *PMR4/GSL5* is expressed ubiquitously in unwounded *Arabidopsis thaliana* plants (Enns et al., 2005). Inoculation with *Plectosphaerella cucumerina* fungi results in a slight increase in *GSL5* expression (García-Andrade et al., 2011). Downy mildew *Hyaloperonospora* and the application of SA and flg22 induce *GSL5* expression (Dong et al., 2008; Keppler et al., 2018). Whether and if so how GSL5 expression or activity is regulated according to the nutrient status remains poorly understood.

Here, we report that callose deposition is induced in root hairs under Pi deficiency, in a manner independent of *PHR1/PHL1* and *LPR1/LPR2*. Our genetic studies revealed that the Pi depletion-induced callose deposition is impaired in *pmr4* (*gsl5*) root hairs.

The loss of *PMR4*-mediated callose deposition is associated with reduced plant growth as well as exaggerated stress symptoms under Pi deficiency, in addition with enhanced powdery mildew resistance. The results indicate *PMR4*-mediated callose deposition in root hairs as an important step in plant adaptation to Pi-limiting conditions.

## Results

### Callose deposition occurs in root hairs under phosphate deficiency, independently of *PHR1/PHL1* and *LPR1/LPR2*

Under low Pi conditions (Pi at 0 or 50 µM), we noticed callose deposition in the epidermal cells, particularly in root hairs of the differentiation zone (Figure 1a, b; O’Lexy et al., 2018). However, callose deposition was not detected in root hairs under the tested N, Fe or Fe/Pi deficient conditions (Figure 1a). The results indicate that the callose deposition is specific to Pi deficiency and is dependent on Fe, consistent with the interplay between Fe and Pi (Müller., 2015; Abel, 2017; Xue et al., 2022). The absence of callose staining in the epidermis of root hair-less *rhd6* mutants (Figure 1c) validated that callose deposition occurs in root hairs under low Pi. In contrast to previous study (Müller et al., 2015), we did not detect callose deposition in the root SCN and cortex cells of the elongation zone under our experimental conditions (Figure 1a, c).

**Figure 1.**
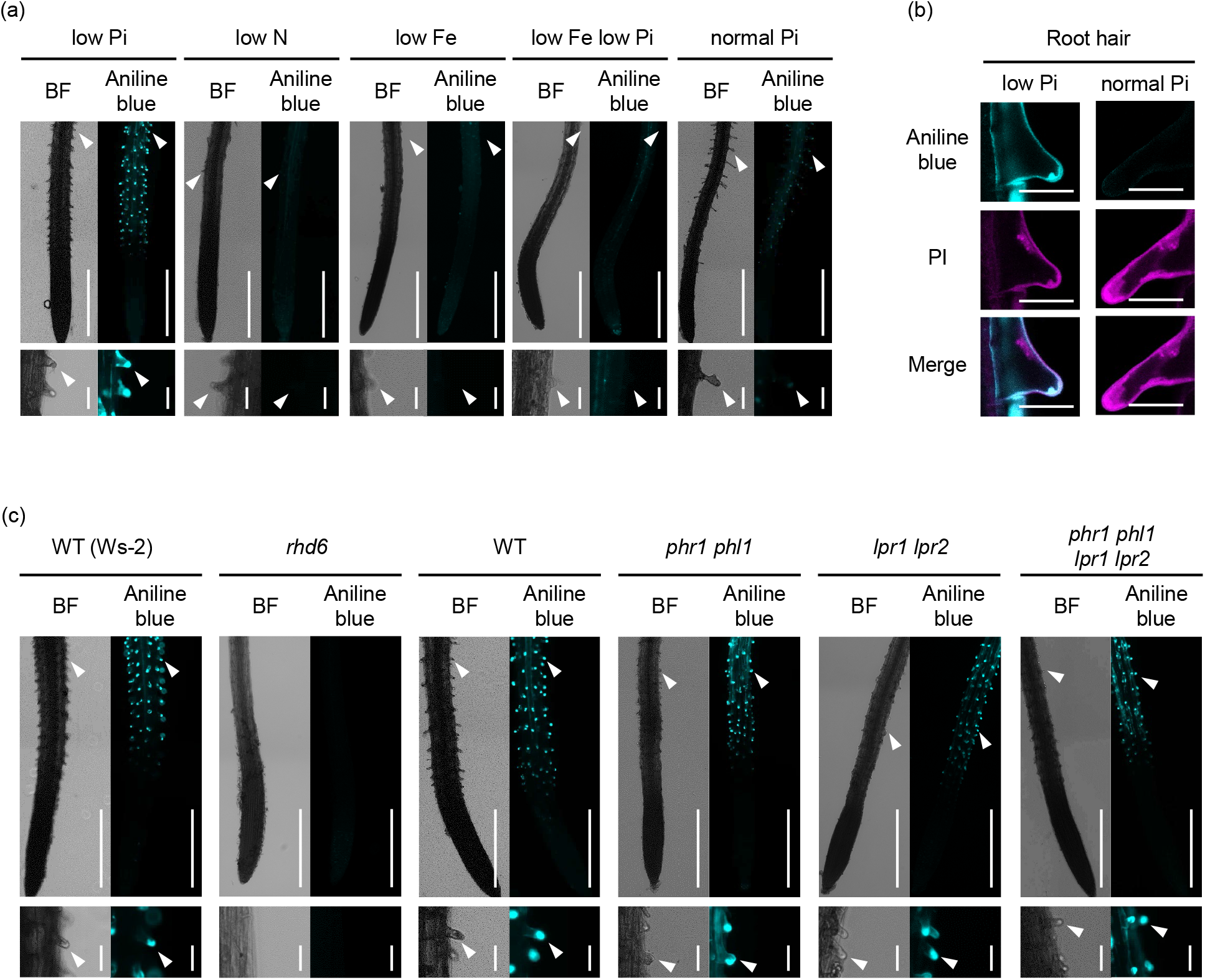
Callose deposition is induced in root hairs under phosphate deficiency, independently of *PHR1, PHL1, LPR1* and *LPR2*. Aniline blue staining of the primary roots 3 d after transfer of 5-d-old *Arabidopsis thaliana* seedlings to low Pi (P 50 µM), low N (N 0 µM), low Fe (Fe 0 µM), low Fe low Pi (Fe 0 µM, P 0 µM) and normal Pi (P 625 µM) liquid media. (a, c, left) Bright field (BF) images. (a, c, right) Aniline blue staining images. White arrowheads indicate root hair tips. (b) Aniline blue and propidium iodide (PI) staining of root hairs under low Pi and normal Pi conditions. (c) The wild-type (WT) used for *rhd6* was Ws-2. Magnified images of the upper panels (a, c, lower panel). Bars, 500 µm (a, c, upper panel), 50 μm (a, c lower panel) and 25 µm (b).

We next assessed whether *PHR1/PHL1* and *LPR1/LPR2* play a role in the root hair callose deposition. Unlike *LPR1*/*LPR2*-mediated callose deposition in SCN (Müller et al., 2015), the root hair callose deposition was not affected in *lpr1 lpr2* (Figure 1c) despite its Fe dependence (Figure 1a). Furthermore, wild type (WT)-like callose deposition was retained in root hairs of *phr1 phl1 lpr1 lpr2* under low Pi conditions (Figure 1c). The results indicate that callose deposition is induced in root hairs under Pi deficiency, independently of these PSR regulators, and its separation from callose deposition in the root SCN.

### *PMR4* callose synthase contributes to callose deposition under Pi deficiency

To unravel genetic requirements for the callose deposition in root hairs under Pi deficiency, we examined *glucan synthase like* (*gsl*) mutants for alternations in callose deposition under phosphate deficiency. Of the tested *gsl* mutants, *gsl5* (hereafter, *pmr4*) mutants specifically lacked callose deposition in root hairs (Figure S1). We also screened a total of 18,550 ethyl methane sulfonate (EMS)-mutagenized individuals in Col-0 and Ler backgrounds. This revealed two mutants in the Col-0 background, designated *<u>c</u>allose <u>a</u>lteration under <u>P</u>i <u>s</u>tarvation 1 and 2* (*caps1 and caps2*, Figure 2a). *caps1* and *caps2* mutants were severely defective in callose deposition of root hairs under low Pi conditions (Figure 2a). DNA Sequencing on the *PMR4* genomic region revealed a DNA substitution within the coding region, resulting in a non-synonymous amino acid substitution, in both *caps1* and *caps*2 alleles (Figure 2b). The substitution in *caps*1 converts Tryptophan to Stop codon at the 260^th^ amino acid (PMR4^W260*^), resulting in PMR4 truncation, consistent with the severely impaired callose deposition in *caps1* as well as *pmr4-1* (PMR4^W687*^). In *caps2*, glutamic acid is substituted with lysine at the 1181^st^ amino acid (PMR4^E1181K^), located within the cytosolic side of a predicted glucan-synthase domain (residue 575 to 1558, orange-marked domain, Figure 2b). The results suggest that dysfunction of PMR4 fails to induce root hair callose deposition under Pi deficiency in these *caps* alleles. We validated that leaf callose deposition in response to the bacterial MAMP flg22, a *PMR4-*dependent output in pattern-triggered immunity (PTI), is abolished in these *caps* alleles (Figure S2).

**Figure 2.**
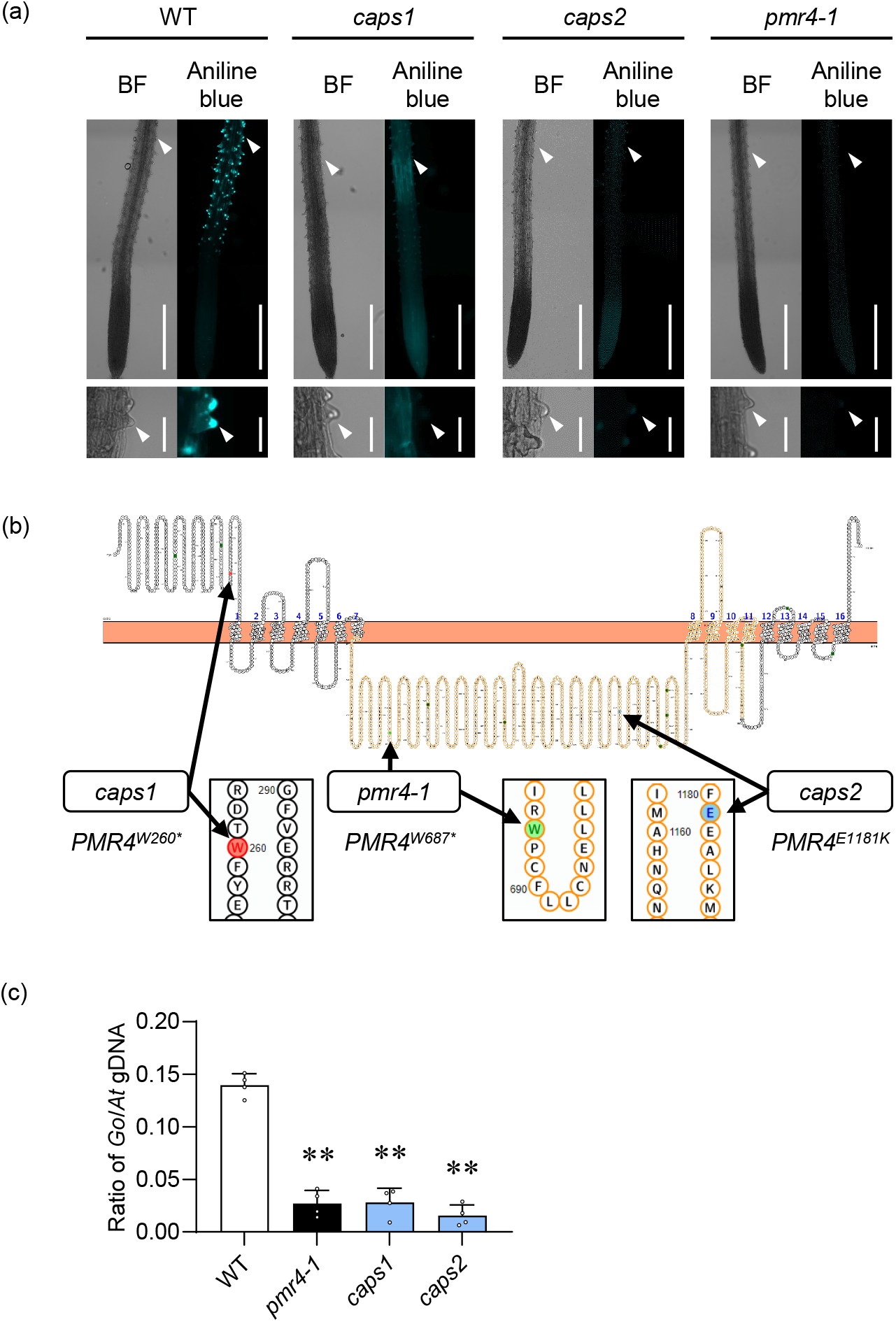
Callose synthase *PMR4* is required for callose induction in root hairs under phosphate deficiency. (a) Aniline blue staining of the primary roots 3 d after transfer of 5-d-old *Arabidopsis thaliana* plants to low Pi (0 µM) liquid media. Left, bright filed (BF) images. Right, aniline blue staining images. Two *caps* mutants are defective in inducing root hair *callose under phosphate starvation.* Lower panels, magnified images of upper panels. White arrowheads indicate root hair tips. Bars, 500 µm (upper panel) and 50 μm (lower panel). (b) Positions of mutated sites in the *PMR4* locus of *caps* mutants, illustrated with Protter (Omasits *et al*., 2014). An orange region corresponds to the catalytic domain of PMR4. Asterisks indicate stop codons. (c) Ratios of *G. orontii* genomic DNA to Arabidopsis genomic DNA were determined by qPCR with the primers for Plasma membrane ATPase 1 (Go_V1_Conting3757) and RNA-binding family protein (AT3G21215), respectively. Data show means ± SD (n=3). Asterisks (** *P* < 0.01) indicate statistically significant differences relative to WT according to Student’s *t*-test.

Moreover, we showed strong resistance to powdery mildew fungi in these *caps* plants (Figure 2c), as previously described in *pmr4* (Vogel & Somerville, 2000; Nishimura et al., 2003). The results confirm that PMR4 functions are impaired in both *caps* alleles.

### *PMR4* influences root development traits induced under Pi deficiency

By using *pmr4-1* as a representative mutant allele, we further investigated the biological significance of *PMR4* under Pi deficiency. Recent studies noted that some of Pi deficiency-induced root responses are limited to the plate-grown plants whose roots are light irradiated (Zheng et al., 2019; Yeh et al., 2020; Gao et al., 2021). To test this possibility, we examined callose deposition in Pi-deplete and Pi-replete plants under darkness. At 3 d after the transfer of seedlings to the different Pi conditions under darkness, we detected callose deposition in non-irradiated root hairs of WT but not *pmr4* plants specifically under low Pi, validating that PMR4-mediated, root hair callose deposition is not a consequence of light exposure to the root (Figure S3a). Moreover, we also detected callose deposition in the cotyledons of WT but not *pmr4* plants under these conditions (Figure S3b). The results strengthen the physiological relevance of PMR4-mediated callose deposition under low Pi.

The loss of *PMR4* leads to an increase in salicylic acid (SA) biosynthesis and defense under biotic stress (Vogel & Somerville, 2000; Nishimura et al., 2003), in a manner requiring *SID2 isochorismate synthase* that mediates a critical step in SA biosynthesis during defense activation (Wildermuth et al., 2001). Given the negative impact of SA defense activation on plant growth (Mateo et al., 2006; Zhang et al., 2010), we assessed the possible SA dependence of the observed *pmr4* phenotypes, by examining genetic interactions between *pmr4* and *sid2*. Callose deposition was induced in *sid2* root hairs but not induced in *pmr4 sid2* under low Pi conditions (Figure 3a), demonstrating that *SID2*-mediated SA biosynthesis is not required for *PMR4*-mediated callose deposition in root hairs. The results also exclude over-activation of SA defense as the cause of the callose defects in *pmr4* root hairs.

**Figure 3.**
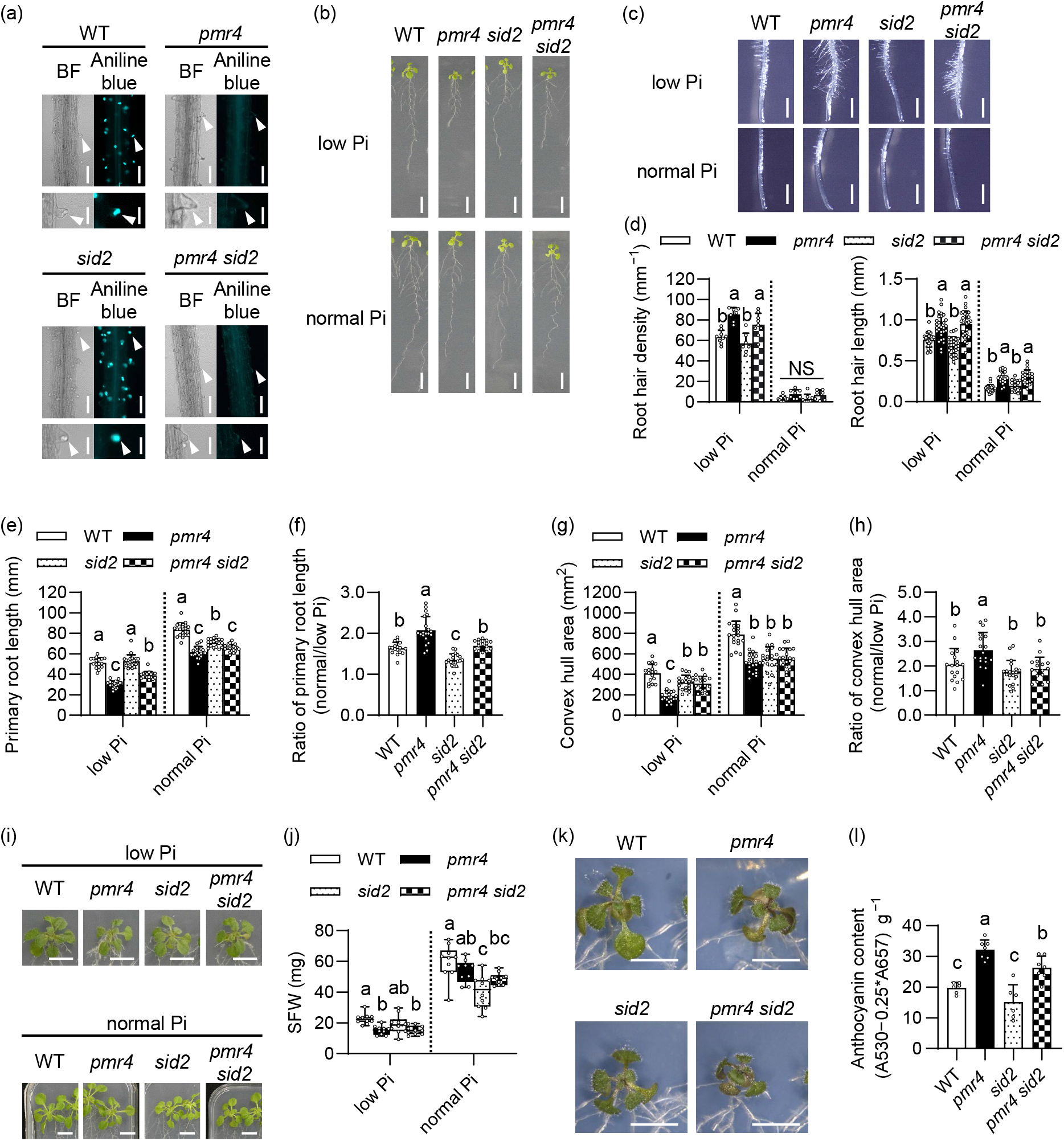
PMR4 modulates root and shoot growth under phosphate deficiency. (a) Aniline blue staining of the primary roots 3 d after transfer of 5-d-old *Arabidopsis* seedlings to low Pi (50 µM) liquid media. Left, bright filed (BF) images. Right, aniline blue staining images. Lower panels, magnified images of upper panels. White arrowheads indicate root hair tips. Bars, 100 µm (upper panel) and 50 μm (lower panel). (b) Six-d-old seedlings were exposed to low Pi (50 µM) or normal Pi (625 µM) media for 6 d. Bar, 10 mm. (c) The primary roots 6 d after transfer of 6-d-old seedling to low Pi (50 µM) and normal Pi media. Bar, 1 mm. (d) Root hair density and root hair length in (c). (e) Primary root length in (b). (f) Ratio of the primary root length under normal Pi relative to low Pi of (e). (g) Convex hull area of the roots in (b). (h) Ratio of convex hull area under normal Pi relative to low Pi of (g). (i) Six-d-old seedlings were exposed to low Pi (50 µM) or normal Pi for 16 d. Bar, 10 mm. (j) SFW in (i). (k) Six-d-old seedlings were exposed to low Pi (10 µM) media for 10 d. Bar, 10 mm. (l) Anthocyanin contents in (k). (d-h, j, l) Data show means ± SD (n = 7∼30). Statistical significance was assessed by one-way ANOVA and Tukey honestly significant different (HSD) test for each medium dataset. Different letters indicate significant differences (*P* < 0.05). NS, not significant.

In Arabidopsis, root hair cells specifically occur at the junction of two underlying cortical cells, whereas non-hair cells occur in the other epidermal cells (Dolan et al., 1994; Schiefelbein et al., 2009). Root hair density was increased under low Pi conditions compared to normal Pi conditions, and the increase was exaggerated in *pmr4* as well as *pmr4 sid2*, without ectopic root hair formation (Figures 3b-d, S4a). Likewise, root hair length was also increased in *pmr4* as well as *pmr4 sid2* under both Pi conditions (Figure 3c, d). The results suggest that *PMR4* is not required for the differentiation or outgrowth of root hair cells, but that *PMR4* contributes to restricting root hair formation and elongation, in the presence or absence of *SID2*. Conversely, primary root length was reduced in *pmr4* compared with WT in both low and normal Pi conditions (Figure 3b, e). We then determined the ratio of the primary root length under normal Pi versus low Pi as an indicator for the primary root growth inhibition under Pi deficiency. The ratio (normal Pi/low Pi) was increased in *pmr4* compared with WT, indicating that primary root growth inhibition, a Pi deficiency-characteristic response, is exaggerated in *pmr4* (Figure 3f). We also observed a decrease in the root meristem region in *pmr4* compared with WT under Pi deficiency, suggesting that the meristem exhaustion coincides with the inhibition of primary root growth in *pmr4* (Figure S4b). The primary root growth inhibition under low Pi was alleviated in *sid2*, but it was again exaggerated in *pmr4 sid2* compared to *sid2* (Figure 3e, f), indicating that the loss of *PMR4* enhances the Pi deficiency response, with or without *SID2*.

We next tested whether and if so how *PMR4* affects the expression of Pi deficiency-inducible genes. Quantitative RT-PCR analyses on WT and *pmr4* roots revealed that the expression of Pi starvation-induced genes, *AT4* and *IPS1* long non-coding RNAs (Franco-Zorrilla et al., 2007), *PHT1;5* Pi transporter (Nagarajan et al., 2011) and *SPX1* Pi-binding suppressor of PHR1 (Puga et al., 2014), was not affected in *pmr4* under the normal (625 µM) and low Pi (50 µM and 10 µM) conditions (Figure S5). The results are consistent with WT-like callose deposition in the root hairs of *phr1 phl1* and *phr1 phl1 lpr1 lpr2* (Figure 1c) and indicate that *PMR4* is not required for the expression of these PSR-related genes.

The primary root growth inhibition under Pi deficiency relies on *LPR1* and *LPR2* (Svistoonoff et al., 2007; Müller et al., 2015). We examined the possible relationship between *LPR1/LPR2* and *PMR4* in the regulation of primary root growth. The primary root length was reduced in *pmr4 lpr1 lpr2* compared with *lpr1 lpr2* under both Pi conditions, whereas it was increased in *pmr4 lpr1 lpr2* compared with *pmr4* specifically under low Pi (Figure S6a, b). The results indicate that *PMR4* serves to sustain the primary root growth irrespective of the Pi availability or *LPR1/LPR2* activity, whereas the *LPR1/LPR2* inhibition is specific under low Pi with or without *PMR4*. The ratio of the primary root length under normal Pi versus low Pi was lowered to a similar extent in *lpr1 lpr2* and *pmr4 lpr1 lpr2* compared with WT (Figure S6c), despite the decrease in the primary root length of *pmr4* (Figure S6b), indicating that *lpr1 lpr2* is epistatic to *pmr4*. Consistently, under Pi deficiency, *PMR4* expression was reduced in *lpr1 lpr2,* but not in *phr1 phl1*, compared to WT roots (Figure S7). Together with the WT-like callose deposition in *lpr1 lpr2* root hairs (Figure 1c), the data suggest that *PMR4* expression is dependent on *LPR1/LPR2* but that the residual *PMR4* expression is sufficient for the callose deposition in *lpr1 lpr2* root hairs.

We next determined the convex hull root area as a proxy of overall root system growth, which was correlated with Pi availability (Figure 3g). It was reduced in *pmr4* and *sid2* compared with WT under both normal and low Pi conditions, albeit to a lesser extent in *sid2* than *pmr4* under low Pi (Figure 3g), suggesting that *PMR4* and *SID2* both contribute to expanding the root area. The ratio of the root area under normal Pi versus low Pi conditions was increased in *pmr4* relative to WT in a manner dependent on *SID2* (Figure 3h), suggesting that *SID2* contributes to low Pi-induced root growth inhibition in *pmr4*. Consistently, the root area was greater in *pmr4 sid2* compared to *pmr4* under low Pi (Figure 3g). The results suggest that SID2 negatively impacts root expansion specifically in the absence of PMR4 under Pi deficiency. The results also imply that SID2-mediated SA biosynthesis promotes root growth expansion in Pi-depleted plants when PMR4 is present. The latter seems to be consistent with a previously described uncoupling of SA induction from defense activation under Pi deficiency (Gulabani et al. 2021).

### *PMR4* serves to sustain shoot growth and restrict anthocyanin accumulation under Pi deficiency

We next examined whether *PMR4* influences shoot growth, by determining shoot fresh weight (FW) under Pi deficiency and sufficiency. Shoot FW was lowered in *pmr4* compared with WT, and it was not even partially restored by the simultaneous loss of *SID2* in *pmr4 sid2* under low-Pi media or soil (Figures 3i-j and S8). Together with the root phenotypes described above, the results exclude *SID2* as the major cause of the growth inhibition in *pmr4*. Although high SA accumulation was previously described to reduce plant growth (Miura et al., 2010), we did not detect negative impacts of *SID2* on shoot growth under low Pi conditions (Figure 3j). *SID2* even positively influenced shoot growth under the normal Pi conditions, indicated by the reduced shoot biomass in *sid2* compared to WT (Figure 3j).

Anthocyanin content was increased in *pmr4* compared with WT, in the presence or absence of *SID2* (Figure 3k, l). Although SA was described to enhance anthocyanin accumulation under Pi deficiency (Morcillo et al., 2019), *sid2* was largely indistinguishable from WT in the anthocyanin contents under our low Pi conditions (Figure 3k, l). The results imply that PMR4-mediated Pi acquisition and allocation serves to alleviate Pi starvation causing anthocyanin accumulation under Pi deficiency.

### *PMR4* promotes the acquisition and root-to-shoot translocation of Pi

The *PMR4* effects on shoot and root growth prompted us to examine its possible role in Pi acquisition from the rhizosphere and Pi mobilization within the plant. We found that shoot Pi concentrations were greatly reduced in *pmr4* compared with WT, without alterations in root Pi concentrations, under Pi deficiency (Figure 4a, b). This indicates that *PMR4* contributes to shoot Pi accumulation in Pi-depleted plants. It was described that *PHR1/PHL1* promotes shoot Pi accumulation (Wang et al., 2018) and that *PHO1* Pi transporter mediates Pi mobilization from roots to shoots through the root xylem vessels (Stefanovic et al., 2007). We validated that Pi contents were lowered in the shoots but increased in the roots, in *phr1 phl1* and *pho1* compared with WT under low Pi (Figure 5a). A decrease in shoot Pi contents was also observed in *phr1 phl1* and *pho1* under normal Pi (Figure 5b). Since root hairs contribute to Pi accumulation under Pi deficiency (Tanaka et al., 2014; Robertson-Albertyn et al., 2017; Holz et al., 2018), we next determined Pi contents in a root hair less mutant, *rhd6*. As seen in *pmr4, phr1 phl1* and *pho1*, Pi contents were lowered in the shoots but not in the roots of *rhd6* compared to WT, specifically under Pi deficiency (Figure 6a, b). As illustrated in Figure 5c and 6c, all these mutants displayed a decrease in shoot Pi contents without reducing root Pi contents under Pi deficiency. We inferred from these results that *PMR4*-mediated callose deposition in root hairs contributes to shoot Pi accumulation, possibly through increasing the uptake or root-to-shoot translocation of Pi. To test the hypothesis, we determined the rates of Pi uptake and Pi translocation by applying and tracing radio-labelled P. The rate of ^32^P uptake per time and the ratio of ^32^P in the shoot to whole seedling, standing for Pi translocation rate, were both lowered in *pmr4* and *pmr4 sid2* compared to WT and *sid2,* respectively, under low Pi (Figure 5d, 6d). In the presence or absence of *SID2*, *PMR4* contribution to the two proxies was significant only under low Pi (Figure 5d, 6d). These results suggest that PMR4 promotes Pi uptake and root-to-shoot Pi translocation under Pi deficiency. Moreover, the ^32^P uptake rate was reduced in *sid2* compared to WT, whereas *pmr4 sid2* was not distinguishable from *pmr4* under low Pi (Figure 6d), suggesting that SID2 positively influences Pi uptake under Pi deficiency in the presence of PMR4.

**Figure 4.**
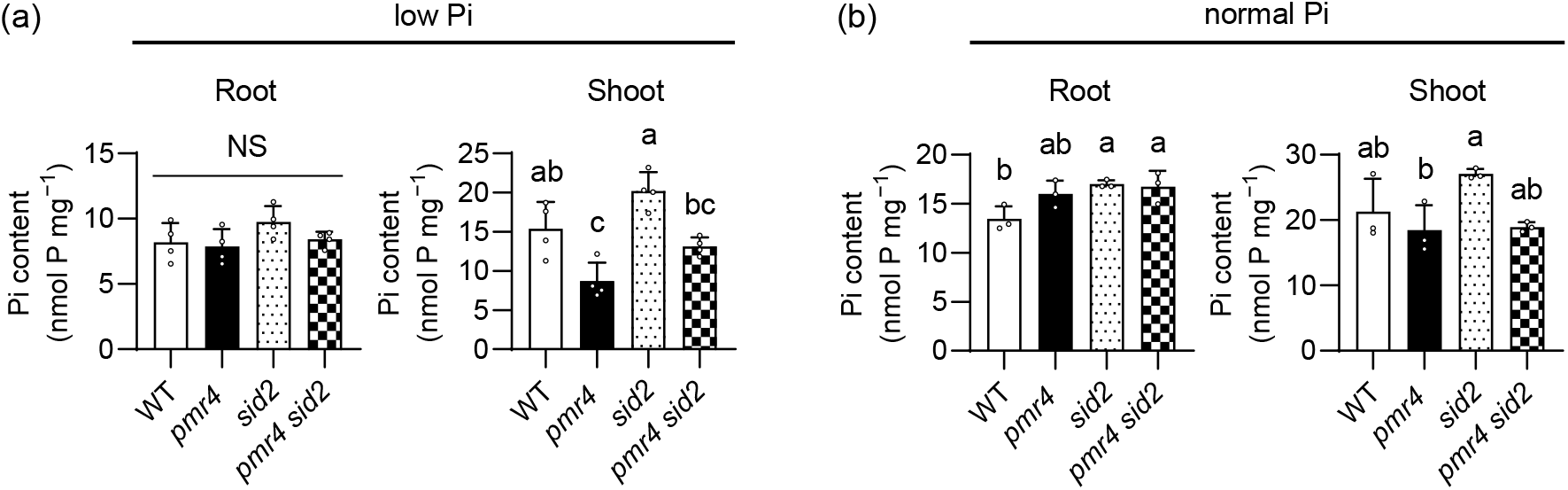
***PMR4* contributes to shoot Pi accumulation under phosphate deficiency.** (a-b) Pi contents per root or shoot fresh weight (FW) of 6-d-old *Arabidopsis* seedlings exposed to low Pi (150 µM) or normal Pi (625 µM) for 9 d. Data show means ± SD (n = 3∼4) in (a-b). Statistical significance was assessed by one-way ANOVA and Tukey HSD test for each medium dataset (a-b). Different letters indicate significant differences (*P* < 0.05). NS, not significant.

**Figure 5.**
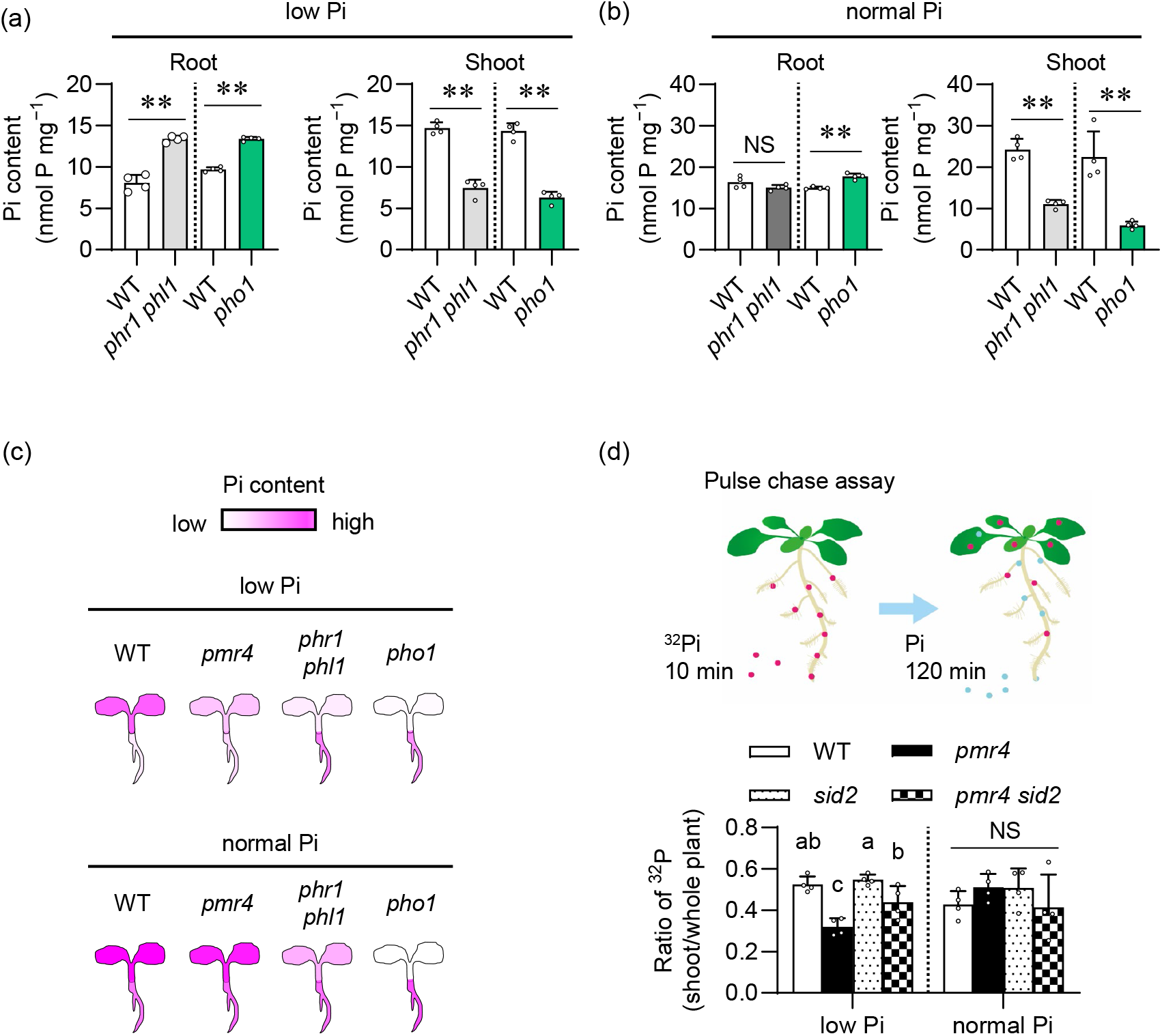
PMR4 contributes to Pi translocation under phosphate deficiency. (a-b) Pi contents per root or shoot fresh weight (FW) of 6-d-old *Arabidopsis* seedlings exposed to low Pi (150 µM) or normal Pi (625 µM) for 9 d. (c) Illustration of Pi accumulation in the indicated genotypes, referred to (Figure 4a, 5a-b). (d) Ratio of ^32^P in the shoots relative to the whole plants after pulse-chase labeling. Seedlings grown on low and normal Pi media (50 and 625 µM, respectively) were exposed to ^32^P-labeled Pi for 10 min, and then transferred to non-labelled Pi media for 120 min (pulse chase). The ^32^P ratio of the shoots to the whole seedlings was calculated for the translocation rate of Pi. Data show means ± SD (n = 3∼4) in (a-b, d). Asterisks (** *P* < 0.01, * *P* < 0.05) indicate statistically significant differences relative to WT according to Student’s *t*-test (a-b). Statistical significance was assessed by one-way ANOVA and Tukey HSD test for each medium dataset (d). Different letters indicate significant differences (*P* < 0.05). NS, not significant.

**Figure 6.**
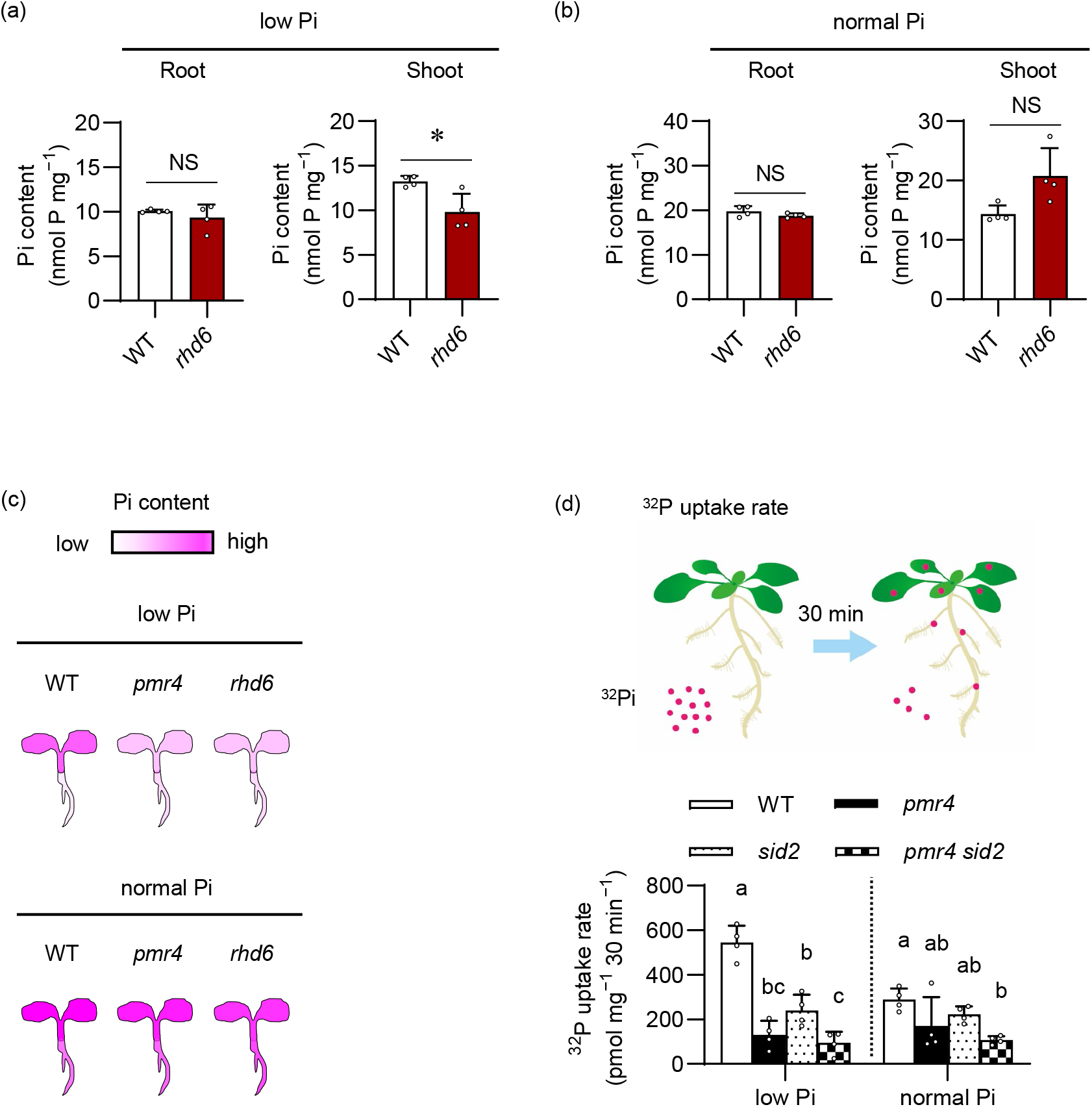
PMR4 contributes to Pi uptake under phosphate deficiency. (a-b) Pi contents per root or shoot fresh weight (FW) of 6-d-old *Arabidopsis* seedlings exposed to low Pi (150 µM) or normal Pi (625 µM) for 9 d. The WT used for *rhd6* was Ws-2 (c) Illustration of Pi accumulation in the indicated genotypes, referred to (Figure 4a, 6a-b). (d) ^32^P uptake rate determined in the whole seedlings exposed to the indicated media (low Pi and normal Pi, 50 and 625 µM Pi, respectively) supplemented with ^32^P-labelled Pi for 30 min. Data show means ± SD (n = 3∼4) in (a-b, d). Asterisks (** *P* < 0.01, * *P* < 0.05) indicate statistically significant differences relative to WT according to Student’s *t*-test (a-b). Statistical significance was assessed by one-way ANOVA and Tukey HSD test for each medium dataset (d). Different letters indicate significant differences (*P* < 0.05). NS, not significant.

We further assessed the possible *PMR4* contributions to the acquisition and tissue mobilization of Pi and other elements, by inductively coupled plasma mass spectrometry (ICP-MS) analyses for P, K, Mg, Ca, Mn, Fe, Zn, Mo and B contents in Pi-depleted and Pi-replete plants. Under Pi sufficiency, although the contents of these elements were not distinguishable between WT and *pmr4* roots, the contents of K and the rest were increased and lowered, respectively, in *pmr4* compared to WT shoots (Figure S9). Under low Pi, Mn, Zn and Mo contents were greater in *pmr4* compared with WT roots, whereas the contents of all the tested elements except K were lowered in *pmr4* compared with WT shoots (Figure 7). The results were consistent with *PMR4*-mediated shoot Pi accumulation without affecting root Pi contents under low Pi conditions (Figures 4a). Additionally, the results suggest that *PMR4* also positively influences shoot accumulation of Mg, Ca, Mn, Fe, Zn, Mo and B (Figures 7, S9).

**Figure 7.**
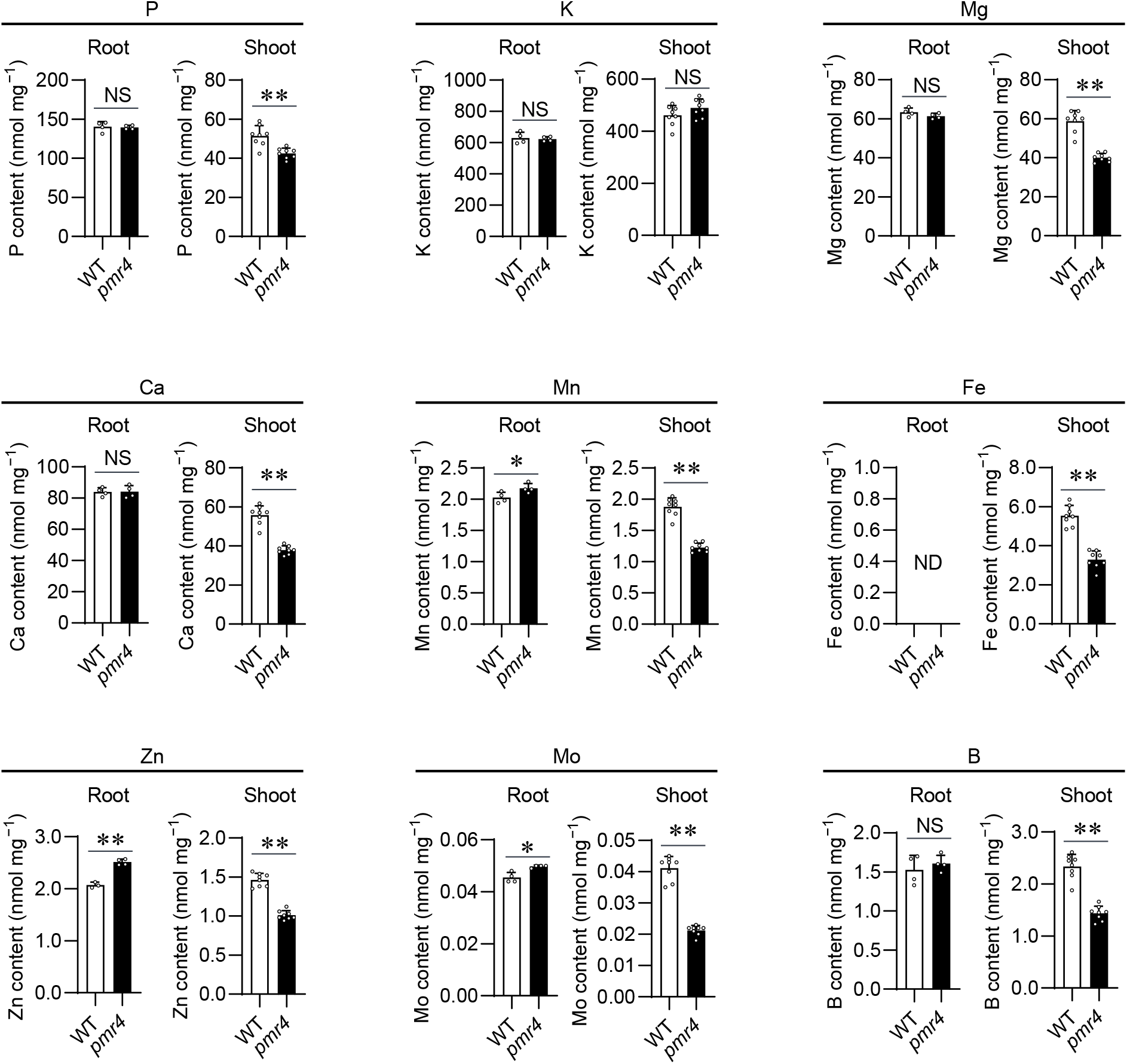
Plant element contents in shoot and root under low phosphate conditions. Element contents (P, K, Mg, Ca, Mn, Fe, Zn, Mo and B) per root or shoot dry weight (DW) of 5-d-old *Arabidopsis thaliana* seedlings exposed to low Pi (10 µM) for 4 d. Data show means ± SD (n=4∼8). Asterisks (** *P* < 0.01, * *P* < 0.05) indicate statistically significant differences relative to WT according to Student’s *t*-test. NS, not significant. ND, no data.

WT-like and increased K accumulation in *pmr4* shoots compared to WT under Pi deficiency and sufficiency, respectively, show that the root-to-shoot transport is not entirely reduced for all elements in *pmr4*, pointing to a certain degree of element specificity in *PMR4* dependence (Figures 7, S9).

The role of *PMR4* in the shoot accumulation of different elements prompted us to examine whether *PMR4* influences plant responses to different nutrient deficiency. We examined *pmr4* phenotypes under B scarcity, as B scarcity induces inhibition of the primary root and transcriptional reprogramming related to ROS and defense in roots (Miwa et al., 2013; Das & Purkait, 2020). Although the primary root length was reduced in *pmr4* compared to WT over the tested range of B conditions, its ratio under B sufficiency (100 µM) versus the examined low B conditions was not affected in *pmr4* (Figure S10). The results point to specific contribution of *PMR4* to plant growth under Pi deficiency.

### Root *PMR4* contributes to shoot growth under Pi deficiency

We next examined the possible significance of the root *PMR4* pool in promoting shoot growth, through micrografting between WT and *pmr4* plants. We validated that the micro-grafted plants of WT/WT (shoot/root genotypes) and *pmr4/pmr4* showed the corresponding WT and *pmr4* phenotypes in root growth and SFW (Figure 8a-c). The root hair callose was detected only when *PMR4* was present in the roots, although the callose signals were not as strong as in intact plants, possibly reflecting the stresses imposed by the grafting procedure (Figure 8d). Remarkably, SFW was greatly reduced in WT*/pmr4* whereas it was less affected in *pmr4/*WT (Figure 8b), pointing to a critical role for a root *PMR4* pool in shoot growth under Pi deficiency. However, the root growth was partially restored in WT/*pmr4* (Figure 8c), pointing to a role for a shoot *PMR4* pool in sustaining the primary root growth in Pi-depleted plants.

**Figure 8.**
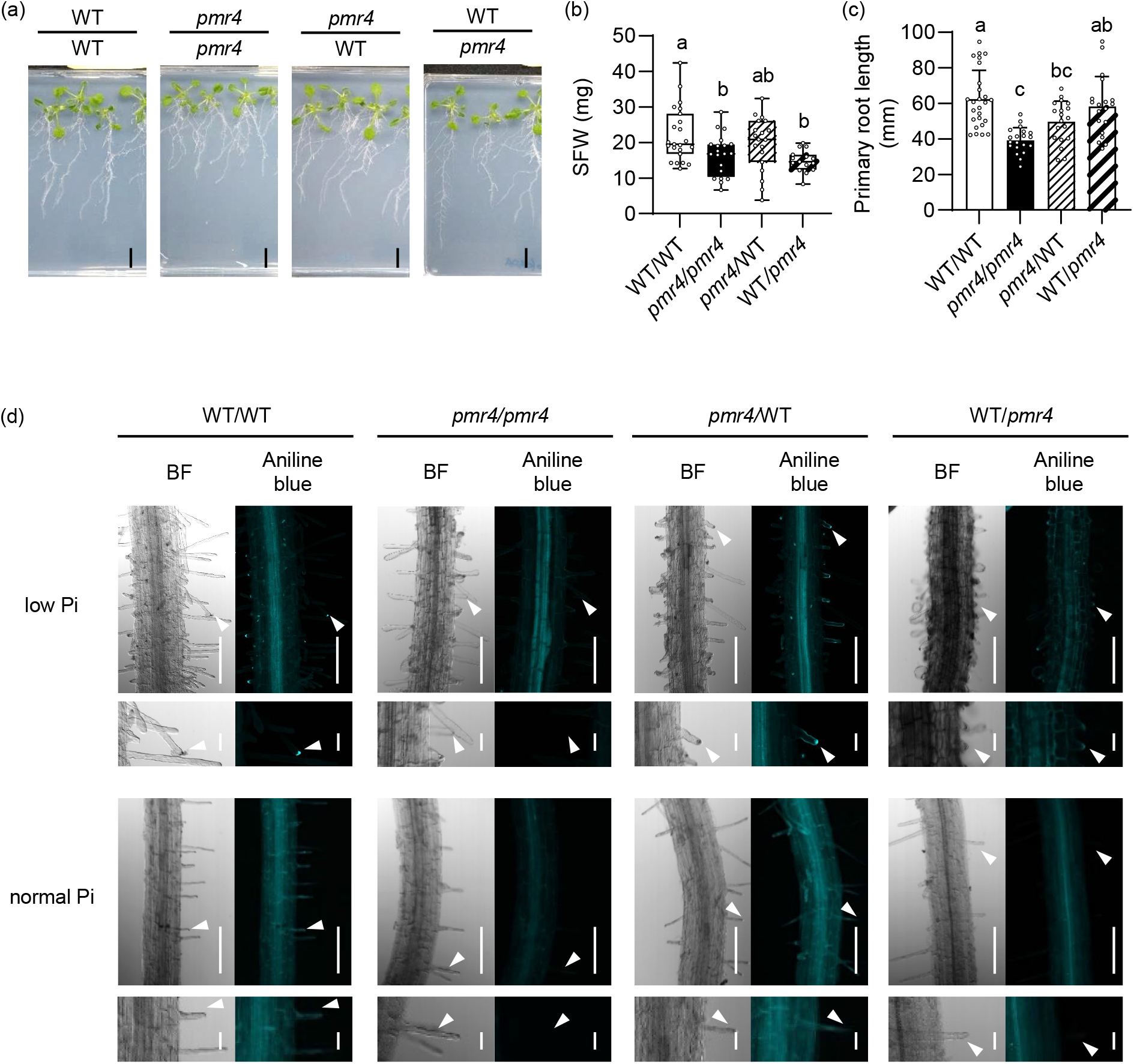
Root PMR4 contributes to shoot growth under phosphate deficiency. (a) Reciprocal micrografting of *Arabidopsis* WT and *pmr4* mutants. Images were obtained at 14 d after transfer to low Pi (150 µM) media. Bar, 10 mm. (b) Shoot fresh weight (SFW) of (a). (c) The primary root length of (a). (d) Aniline blue staining of grafted-seedlings at 9 d after transfer to low Pi (50 µM). (d, left) Bright filed (BF) images. (d, right) Aniline blue staining images. (d, lower panel) Magnified images of (d, upper panel). White arrows indicate root hair tips. Bar, 100 µm (c, upper panel) or 20 μm (c, lower panel). (b, c) Data show means ± SD (n=19-26). Statistical significance was assessed by one-way ANOVA and Tukey HSD test for each medium dataset. Different letters indicate significant differences (*P* < 0.05).

To further examine the role of a PMR4 pool in root hairs, we generated transgenic plants expressing *PMR4* under the control of a root hair-specific *LRX1* promoter (*pLRX1*, Baumberger et al., 2003) in the *pmr4-1* background (Figure 9a, b). *pLRX1::PMR4* complemented *pmr4* in terms of the callose deposition in root hairs and shoot growth under low Pi conditions (Figure 9a-c). By contrast, the primary root growth remained suppressed in the *pLRX1::PMR4* transgenic lines as well as in *pmr4* (Figure 9b, d). The results indicate that PMR4 in root hairs is sufficient to restore shoot growth, but not the primary root growth, under Pi deficiency.

**Figure 9.**
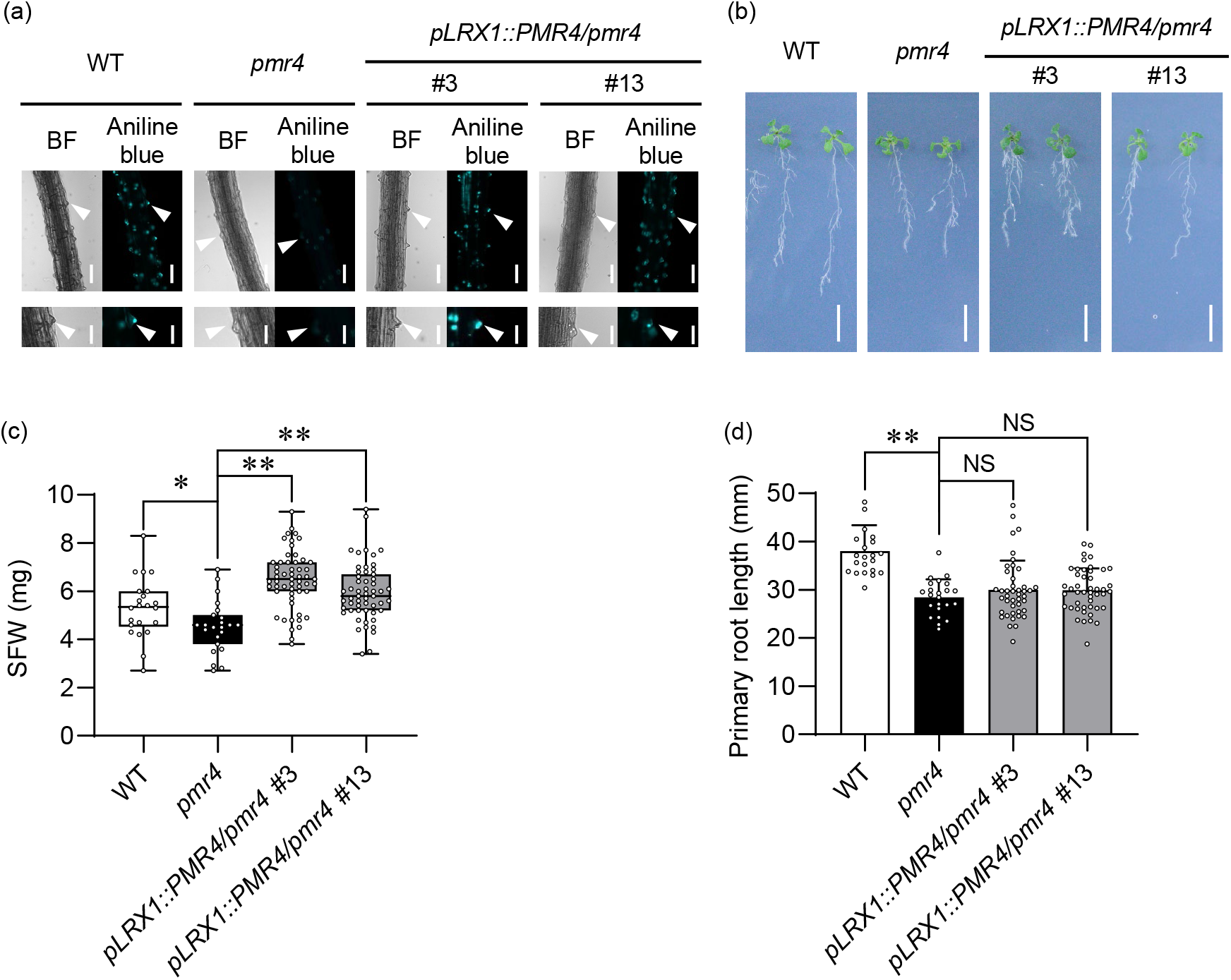
PMR4 in root hairs contributes to shoot growth under phosphate deficiency. (a) Aniline blue staining of the primary roots 3 d after transfer of 5-d-old Arabidopsis seedlings to low Pi (10 µM). White arrowheads indicate root hair chips. Bars, 100 μm (upper panel) and 50 μm (lower panel). (b) Five-d-old seedlings were exposed to low Pi (50 µM) for 10 d. Bar, 10 mm. (c) SFW of (b). (d) Primary root length of (b). (c, d) Data show means ± SD (n = 22-48). Asterisks (** P < 0.01, * P < 0.05) indicate statistically significant differences relative to pmr4 according to Student’s t-test. NS, not significant.

## Discussion

Our studies on callose deposition in root hairs induced under Pi deficiency revealed *PMR4* as a positive regulator of plant adaptation to Pi deficiency (Figure 10). *PMR4* mediates defense-induced callose deposition in pattern triggered immunity and pre-invasion fungal resistance (Jacobs et al., 2003; Chowdhury et al., 2014), and that its loss is linked to enhanced SA-based pathogen resistance (Nishimura et al., 2003). Therefore, *PMR4* seems to play a critical role in an array of biotic and abiotic stress responses, possibly through callose deposition at different sites.

**Figure 10.**
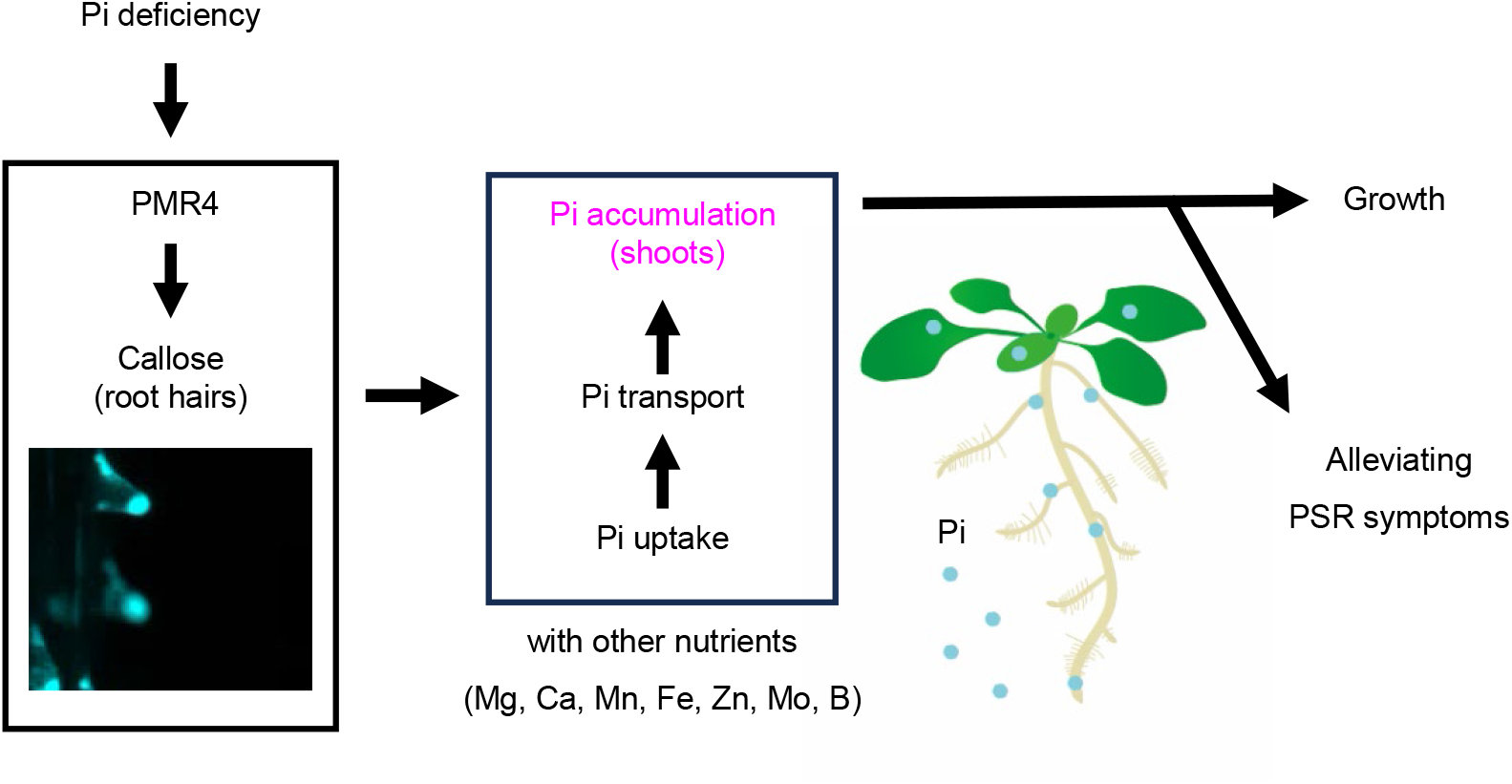
Model for PMR4-mediated root hair callose deposition in plant adaptation to Pi deficiency. PMR4 induces callose deposition in root hairs under phosphate deficiency, and thus contributes to shoot Pi accumulation and growth through enhancing Pi uptake and root-to-shoot transfer. This contribution alleviates phosphate starvation symptoms, such as root system area reduction and anthocyanin accumulation.

The present study suggests that the root hair callose contributes to root Pi uptake and shoot Pi accumulation under Pi deficiency. Importantly, the loss of *PMR4* affects both *PHR1*-dependent and *LPR1*-dependent processes (Figures 3, 4) (Bates & Lynch 1996; Thibaud et al., 2010), which have been genetically uncoupled in PSR (Wang et al., 2010; Balzergue et al., 2017). Pi accumulation and anthocyanin accumulation are suppressed in *phr1 phl1* (Rubio et al., 2001; Bustos et al., 2010; Wang et al., 2018).

Shoot Pi contents and plant biomass are also reduced in *pmr4*, implying that *PMR4* positively influences *PHR1/PHL1*-mediated Pi acquisition and plant growth under internal Pi deficiency. However, over-accumulation of anthocyanin in *pmr4* (Figure 3k, l) in contrast to *phr1 phl1* (Rubio et al., 2001; Bustos et al., 2010; Wang et al., 2018), suggests that *PMR4* contribution is specific to the former two outputs of *PHR1/PHL1* and that its loss results in exaggeration of Pi starvation symptom. The loss of *PMR4* also strengthens inhibition of the primary root growth under Pi deficiency, which is pronounced in the presence of *LPR1/LPR2* (Figure S6), consistent with *LPR1/LPR2-*-dependent *PMR4* expression (Figure S7). The LPR1 pathway mediates callose deposition at the root SCN to inhibit the cell-to-cell movement of the transcription factor SHR, thereby suppressing the cell division and subsequent root elongation (Müller et al., 2015; Mora-Macías et al., 2017). Our results suggest a role for PMR4 in antagonizing LPR1/LPR2-dependent root growth inhibition.

The failure of a root hair PMR4 pool to restore primary root growth (Figure 9b, d) implies the involvement of PMR4 from another cell type(s) than root hairs in this regulation. Consistently, shoot Pi accumulation is partially and essentially retained in *rhd6* under low and normal Pi conditions, respectively, despite the lack of root hairs (Figure 6a-c). It is conceivable that PMR4-mediated callose deposition in different cell types serves to sustain the primary root growth against LPR1/LPR2-mediated attenuation under Pi deficiency. In line with this, we observed low Pi-induced, PMR4-mediated callose deposition in cotyledons as well under darkness (Figure S3). The mechanisms underlying antagonistic interactions between *PMR4* and *LPR1/LPR2* require further studies including the full identification of critical callose deposition sites.

Shoot Pi accumulation depends on root Pi uptake and subsequent Pi translocation. *PHT1;1* and *PHT1;4* provide two major Pi transporters in roots, contributing greatly to Pi acquisition under Pi deficiency (Shin et al., 2004). *PHO1* mediates Pi translocation from roots to shoots through Pi loading into the root xylem (Poirier et al., 1991; Hamburger et al., 2002; Liu et al., 2012). Notably, the loss of *PMR4* results in a decrease in the shoot Pi contents without affecting the root Pi contents (Figures 4a-b, 7). Likewise, the loss of *PHR1/PHL1* and *PHO1* also lowers shoot Pi contents without reducing root Pi contents (Figure 5a, b; Liu et al., 2012; Wang et al., 2018). In *pht1;1 pht1;4*, a decrease of Pi uptake in roots is associated with that of Pi translocation to the shoots and shoot Pi accumulation (Shin et al., 2004). An intimate linkage between root uptake and root-to-shoot translocation is also seen for K^+^ (Nieves-Cordones et al., 2019). It is thus conceivable that the lowered Pi accumulation in *pmr4* shoots is caused by a decrease in the rates of the root Pi uptake and the root-to-shoot Pi translocation, possibly through the compromised root hair function in the absence of *PMR4*. Indeed, ^32^P tracer analyses indicate that *PMR4* promotes both root Pi uptake and root-to-shoot Pi translocation, without affecting the root Pi contents (Figures 4, 5d, 6d). This is consistent with the view that the root-to-shoot Pi translocation is driven by the Pi uptake in the roots (Shin et al., 2004). Leaf transpiration affects root-to-shoot solute translocation (Nieves-Cordones et al., 2019), but it might be ignorable under constitutive high humidity in the closed plate assays used in the present study. Notably, *PMR4* also contributes to the accumulation of different elements except K in shoots under Pi deficiency (Figures 7, S9). This might largely attribute to secondary effects of *PMR4*-dependent root growth and expansion (Figure 3e-h), but the separate regulation of K (Figures 7, S9) cannot be explained by this model. Notably, the disruption of multiple *PHT1* members (*PHT1;1/1;2/1;3* silencing in the *phf1 pht1;4* mutant background, designated mut5) results in a decrease in Na as well as P contents compared to WT (Ayadi et al., 2015). *PHR1* mediates cross-regulation among Fe, Zn and Pi homeostasis (Briat et al., 2015). Further studies will be required to unravel the mechanisms underlying the complex coordination of the acquisition and tissue distribution among different elements and the observed role of PMR4 in the regulation (Figures 7, S9).

Root hairs increase the root surface area to promote root anchorage, nutrient acquisition, and metabolite exudation, thereby facilitating the interactions with the soil environments (Datta et al., 2011; Poole et al., 2018; Vissenberg et al., 2020; Robe et al., 2021; Han et al., 2023). The elongation and increased densities of root hairs in *pmr4* (Figure 3c, d) can be inferred that the root hairs formed in the absence of callose deposition are less functional in Pi acquisition or Pi allocation to the other tissues, thereby leading to further formation of root hairs.

How does PMR4-mediated callose deposition contribute to the root hair functions under low Pi? Our findings are reminiscent of the nutrient transport through the phloem that requires callose deposition in sieve plates (Barratt et al., 2011, Xie & Hong, 2011). The loss of callose from sieve plate pores reduces the phloem conductivity and thus the phloem transport (Barratt et al., 2011). Therefore, it is tempting to speculate that the root hair callose plays a role in cell wall remodeling to reinforce the root hairs for effective transport of water and nutrients and for providing a driving force for their root-to-shoot transport. The precise mechanisms by which callose deposited in the root hairs promotes Pi acquisition require further investigation.

Root hairs also provide a key interface to soil-borne microbes and are intimately associated with root defense regulation. Root hair formation and elongation are facilitated by jasmonate (JA) (Zhu et al., 2006), which mediates the expression of key transcription factors for root hair differentiation and the de-repression of RHD6 from the interference by JAZ proteins (Han et al., 2020). In beneficial interactions with the epiphytic bacterium *Pseudomonas simiae* WCS417, activation of defense-related genes is increased in hair cell lineages compared to non-hair cell lineages (Verbon et al., 2023). The endogenous elicitor-active Pep peptides induce root hair formation (Okada et al., 2021; Jing et al., 2024), and Pep receptor signaling from the root hairs can activate defense responses without root growth inhibition (Okada et al., 2021). These findings suggest a specific capacity of root hair cells in root immunity. How these properties of root hairs are influenced by *PMR4*-mediated callose deposition under Pi deficiency also represents an interesting topic for future studies.

## Experimental procedures

### Plant materials and growth conditions

The *Arabidopsis thaliana* accessions used were Col-0 and Ws-2, including the mutants examined (Table S1) and the wild type used was Col-0 unless otherwise stated. We generated ethyl methanesulfonate (EMS)-mutagenesis (M2) seeds for screening in both Col-0 and Ler backgrounds. All seeds were sterilized using 6% sodium hypochlorite and 0.2% Trion X-100 for 15 min, and then kept at 4°C for 2 d before sowing a half-strength Murashige and Skoog (MS) medium with 25 mM sucrose and 0.8% Agar (WAKO, Japan). If not stated otherwise, the normal Pi media contained 625 μM KH_2_PO_4_, whereas the low Pi media contained 0 µM, 10 µM, 50 μM or 150 μM KH_2_PO_4_ as indicated. The low N and low Fe media used contained 0 µM N and 0 µM Fe, respectively. Plants were grown in a growth chamber under 12 h: 12 h, white light: dark cycles at 22°C unless otherwise stated. The white light intensity was 80 μmol m^−2^ s^−1^. Six-day (d)-old seedlings were transferred to the indicated Pi liquid media or agar media with 1% Difco Agar, Bacteriological (BD, Franklin Lakes, NJ, USA).

### Construction and plant transformation

The genomic DNA sequences of the *POWDERY MILDEW RESISTANCE 4* (*PMR4*, AT4G03550) loci, together with a regulatory DNA sequence for root hair specific expression (below), were cloned into the Gateway binary vector pGWB504 (Nakagawa et al., 2007; primer information Table S2) to express a C-terminally GFP-fused PMR4 under its native regulatory DNA sequence. The root hair-specific promoter sequence used was derived from *LRR-EXTENSIN 1* (*LRX1*, AT1G12040) (Baumberger et al., 2003). Transgenic lines were generated by Agrobacterium-mediated transformation of *pmr4-1* with the DNA construct and subsequently selected on hygromycin resistance and DNA genotyping.

### Aniline blue and PI staining

Callose staining with aniline blue was conducted as described in Schenk & Schikora (2015). Plants were examined at 3 d after the transfer of 5-d-old seedlings to the indicated Pi, N or Fe liquid media (Figure 1a, c). Plant tissues were fixed in a solution containing acetic acid and ethanol (1:3 v/v) overnight. The fixation solution was exchanged to a 150 mM K_2_HPO_4_ solution before callose staining with 0.1% aniline blue in the K_2_HPO_4_ solution for 2 h. The data presented are representative of three independent experiments (Figures 1a, 1c, 2a, 3a), two independent experiments (Figures 1b, 6d, S1, S2) and one experiment (Figures 7a, S3). To localize the root meristematic regions, the primary roots were stained with 20 μg/ml propidium iodide (PI). Callose or PI signals were examined under microscopy using a fluorescence microscope DMI 6000B (Leica Microsystems, Wetzlar, Germany) and a confocal microscope SP8 (Leica Microsystems, Wetzlar, Germany).

### Quantitative real-time PCR analysis

Three-day-old seedlings were exposed to low Pi or normal Pi media for 4 d. Total RNA was then extracted from the root tissues by NucleoSpin® RNA Plant kit (Machete-Nagel, Dueren, Germany). Total RNA was reverse-transcribed using the PrimeScript RT reagent kit including a genomic DNA eraser (Perfect Real Time, Takara Bio, Ohtsu, Japan). Quantitative PCR was performed with SYBR Green (Life Technologies, Carlsbad, CA, USA) and the Thermal Cycler Dice® Real-Time System (TakaraBio). Expression levels of the examined genes (Table S2) were normalized relative to a reference gene, *ACTIN 2* (*ACT2*). PSR-inducible markers examined were *AT4* (At5g03545), *IPS1* (AT3g09922), *PHT1;5* (AT2g32830) and *SPX1* (AT5g20150). Data are representative of two independent experiments

### Identification of mutants altered in the root hair callose deposition

We examined callose deposition in *gsl* mutants (Table S1) after aniline blue staining of the roots transferred to the low Pi liquid media (50 μM) for 3 d. In the EMS mutagenic screening, we subjected a region ∼5 cm from the root tip to aniline blue staining, after cutting out of the M2 individuals transferred to the Pi -depleted liquid media (0 μM) for 3 d. This enabled to keep the plant alive for subsequent characterization and seed propagation. The *PMR4* genomic region was sequenced in the selected mutant candidates.

### Inoculation analyses with powdery mildew fungi

Inoculation of *Golovinomyces orontii* and determination of fungal versus plant DNA ratio were conducted essentially by following Weßling & Panstruga (2012).

Eighteen-d-old plants grown in 10 h: 14 h, light: dark cycles at 22°C were inoculated with *G. orontii* spores at approximately 750 spores cm^−2^. The inoculated leaves were harvested and frozen in liquid nitrogen before determining fungal biomass at 10 d after inoculation. Genomic DNA was extracted essentially by following Brouwer et al. (2001). Quantitative real-time PCR was performed with primers (Table S2) and Brilliant Sybr Green QPCR Reagent Kit (Stratagene, Waldbronn, Germany). The ratio of *G. orontii* to Arabidopsis genomic DNA was calculated using the ΔΔCt method (Pfaffl, 2001). Data are representative of two independent experiments (Figure 2c).

### Determination of root hair density, primary root length and root area

Six-day-old seedlings were transferred and further grown in the low (50 µM) or normal Pi media (625 µM) for 6 d, and then root hairs, primary root and root system area were examined under stereomicroscope M80 (Leica Microsystems, Wetzlar, Germany). The number of root hairs was determined in a region 2.0-3.0 mm from the root tip using Image J software (National Institutes of Health, Bethesda, MD, USA). Data are representative of similar results from three independent experiments (Figure 3c, d) and two independent experiments (Figure S4). The primary root length and convex hull area were determined using ImageJ software and Root nav software (Pound et al., 2013).

Data are representative of three independent experiments (Figure. 3e-h). For growth under different Boron (B) conditions, plants were grown in MGRL solid media containing 1% sucrose and 1% gellan gum (WAKO, Japan). B concentrations were adjusted with boric acid. Plants were grown under 16 h: 8 h, light: dark cycles at 22°C. The data presented are from one independent experiment (Figure S10).

### Determination of shoot fresh weight, anthocyanin and Pi contents

In plant growth assays on media, six-day-old seedlings were transferred to the indicated media (10 μM, 50 μM or 625 μM Pi) and then further grown for the indicated days.

Shoot fresh weight (SFW) was determined 16 d after transfer to the indicated media (50 μM or 625 μM Pi; Figure 3i, j). In plant growth assays with *pLRX1::PMR4* lines, SFW was determined 10 d after the transfer to the low Pi media (10 μM :Figure 9a, c). In plant growth assay on soil, organic black soil (Maruishi Engei, Osaka, Japan) and Ezo sand (Kobayashi Sangyo, Kita-Nagoya, Japan) mixed at 1:1 (v/v) were used as low Pi soil, and Sakata-Super-Mix-A (Sakata Seed, Kyoto, Japan) and Vermiculite G20 (Maruishi Engei, Osaka, Japan) mixed at 1:1 (v/v) were used as normal Pi soil (Figure S8). SFW of seedlings grown on soil was determined 14 d after the transfer of 10-d-old seedlings to the low and normal Pi soils. Data are representative of two independent experiments (Figures 3i, 3j, S5) and one independent experiment (Figure 9b, c).

In the determination of anthocyanin content, six-day-old seedlings were transferred to and further grown on low Pi media (10 μM) for 10 d (Figure 3k, l). The shoots were subject to the procedure described in Mita et al. (1997). In brief, a HCl: methanol mixture at 1:99 (v/v) was added to the ground tissues with 1 ml per sample, and then the resultant homogenates were incubated at 4°C overnight. *A*_530_ and *A*_657_ of the supernatants were determined with a spectrometer UV-1800 (Shimadzu, Kyoto, Japan). Anthocyanin content was calculated following the formula: (A_530_ − 0.25 × A_657_) × fresh weight (FW)^−1^. Data are representative of two independent experiments with the same conclusions (Figure 3k, l).

In the determination of Pi content, six-d-old seedlings were transferred to and further grown on the low Pi (150 μM) or normal Pi (625 μM) media for 9 d (Figures 4, 5a-b, 6a-b). Pi content was determined by following Ames (1966). In brief, the extraction buffer (10 mM Tris, 1 mM EDTA, 100 mM NaCl, 1 mM phenylmethylsulfonyl fluoride pH8.0) was added with 10 µl per sample mg FW. The homogenate 100 µl was mixed with 1% glacial acetic acid 900 µl, and then incubated at 42°C for 30 min. The supernatant was recovered to be incubated in a solution (0.35% NH_4_MoO_4_, 0.86 N H_2_SO_4_, and 1.4% ascorbic acid) at 42°C for 30 min. *A_820_* of the aliquot was measured under Eppendorf BioSpectrometer® basic (Eppendorf, Hamburg, Germany). Then, Pi content was calculated by a P calibration curve prepared with a P standard solution.

Data are representative of three independent experiments (Figure 4a, b) and two independent experiments (Figures 5d, 6d) with the same conclusions.

### Scanning electron microscope

The maturation zone of primary roots 10 d after transfer of 5-d-old seedlings to low Pi (50 µM) liquid media was fixed in half-Karnovsky’s fixative overnight at 4°C, and then postfixed in 1% osmium tetroxide for 1 h at room temperatures (Karnovsky, 1965). The specimens were dehydrated in an ethanol series (30%, 50%, 70%, 80%, and 90%, and 100% at three times) for 10 min each at RT, and then the specimens were dipped in t-butyl alcohol and dried with a vacuum evaporator. They were coated with vapor of 1% osmium tetroxide and then observed with a scanning electron microscope (JIB-4601F, JEOL, Tokyo, Japan).

### Inductively coupled plasma-mass spectrometer (ICP-MS) analyses

Five-day-old seedlings were transferred to and further grown on the growth media (10 μM or 625 μM KH_2_PO_4_) for 4 d (Figures 7, S9). Shoots and roots were harvested and dried at 60°C. After determining the dry weight, the dried tissues were digested with 2 ml nitric acid and 0.5 ml hydrogen peroxide (Wako Pure Chemicals) at 110°C. Digested samples were then dissociated with 5 ml of 2% (w/v) nitric acid and subjected to inductively coupled plasma mass spectrometry (NexION 2000, Perkin-Elmer). The contents of P, K, Mg, Ca, Mn, Fe, Zn, Mo and B were determined. The data were obtained from one experiment (Figures 7, S9).

### Determination of ^32^P uptake rate and translocation ratio

Seven-day-old seedlings were transferred to and further grown on the growth media (50 or 625 μM KH_2_PO_4_) for 10 d. The seedlings were exposed to the liquid media containing ^32^P-labeled Pi (^32^PO_4_ ^3-^ 20 kBq / ml) for 30 min. The plant tissues were fixed to an Imaging Plate (GE Healthcare co., Chicago, USA) for 3 h and then scanned by Typhoon FLA 9000 (GE Healthcare co., Chicago, USA). ^32^P uptake rate was calculated based on a calibration curve for ^32^P. For determining the Pi translocation rate into shoots, the seedlings were exposed to the Pi-containing liquid media containing ^32^P-labeled Pi for 10 min, and then transferred to non-labelled agar media for 120 min. The translocation rate of Pi was determined by calculating ^32^P ratio of the shoot to the whole seedlings. The data presented are representative of two independent experiments with the same conclusions (Figures 5d, 6d).

### Grafting experiments

Grafting of WT and *pmr4* was performed with a supportive micro-device as described in Tsutsui et al. (2020). Four-day-old seedlings grown on a 1/2 MS medium containing 0.5% sucrose and 1% agar under constant light at 22°C were subject to grafting. Both hypocotyls of stock and scion were cut horizontally with a knife and assembled on the device using forceps. The grafted plants were grown in a 1/2 MS medium containing 0.5% sucrose and 2% agar at 27°C for 6 d. In callose staining, the grafted plants transferred and grown on low Pi media (50 µM) under constant light at 22°C for 9 were examined. In plant growth assays, the grafted plants transferred and grown on low Pi media (150 µM) under constant light at 22°C for 14 d were examined. The data presented are representative of two independent experiments with the same conclusions (Figure 8).

## Acknowledgements

We thank the Arabidopsis Biological Research Centre for published plant materials, Yaichi Kawakatsu for advice on micrografting, and Michiko Tsukamoto and Mie Matsubara for technical assistance. This work was supported in part by the grants from Mitsubishi Foundation (YS), the Japan Science and Technology Agency (JST, JPMJPR13B6 and JPMJSC1702 (YS) and JPMJPR16Q7 (KH)), and MEXT of Japan 18H02467 and 21H02507 (YS).

## Author contributions

YS conceived the study. KO and YS designed the experiments. KO, KY, TANN, SK, SY, HT, CT, THL, UN, KI, NT, TI, KM, TM and MN developed the methods, performed the experiments, and/or analyzed the data. SM, MN and KH advised on the experiments. KO and YS wrote the manuscript with contributions from the other authors.

## Supporting Information

Figure S1. Callose deposition in gsl mutants under phosphate deficiency.

Figure S2. flg22-induced callose deposition is impaired in the caps and pmr4 alleles. Figure S3. Pi deficiency-induced callose deposition under dark conditions.

Figure S4. Root hair and meristem morphology in pmr4 under Pi deficiency.

Figure S5. PSR marker gene expression in pmr4 roots

Figure S6. PMR4 restricts LPR1/LPR2-mediated inhibition of the primary root growth under low Pi.

Figure S7. PMR4 expression in phr1 phl1 and lpr1 lpr2 roots. Figure S8. Plant growth is lowered in pmr4 on soil.

Figure S9. Plant element contents in shoot and root under phosphate sufficiency. Figure S10. Primary root growth inhibition under B deficiency.

Table S1. Arabidopsis thaliana mutants used in this study.

Table S2. DNA Primers used in DNA vector construction and qPCR analyses.

