## Supplemental information for "Defense-related callose synthase *PMR4* promotes root hair callose deposition and adaptation to phosphate deficiency in *Arabidopsis thaliana*"

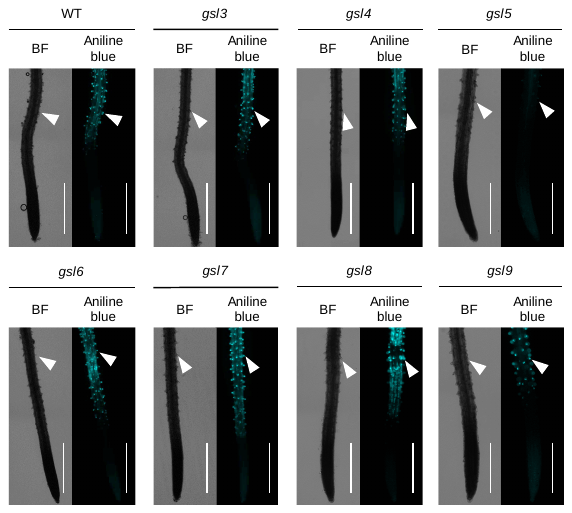

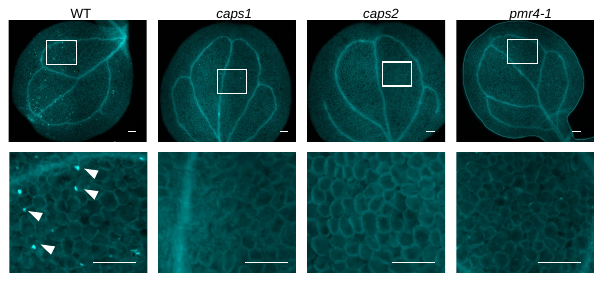

**Figure S1. Callose deposition in *gsl* mutants under phosphate deficiency.**

Aniline blue staining of primary roots 3 d after transfer of 5-d-old *Arabidopsis thaliana* seedlings to low Pi (50 µM) liquid media. Left, bright field (BF) images; Right, aniline blue staining images. White arrowheads indicate root hair tips. Bar, 500 μm.

**Figure S2. flg22-induced callose deposition is impaired in the *caps* and *pmr4* alleles.**

Five-d-old *Arabidopsis thaliana* seedlings exposed to 1 μM flg22 for 24 h were subjected to aniline blue staining. Magnified images of the square area in the upper panels are shown in lower panels. White arrowheads indicate callose deposition. Bars, 200 µm.

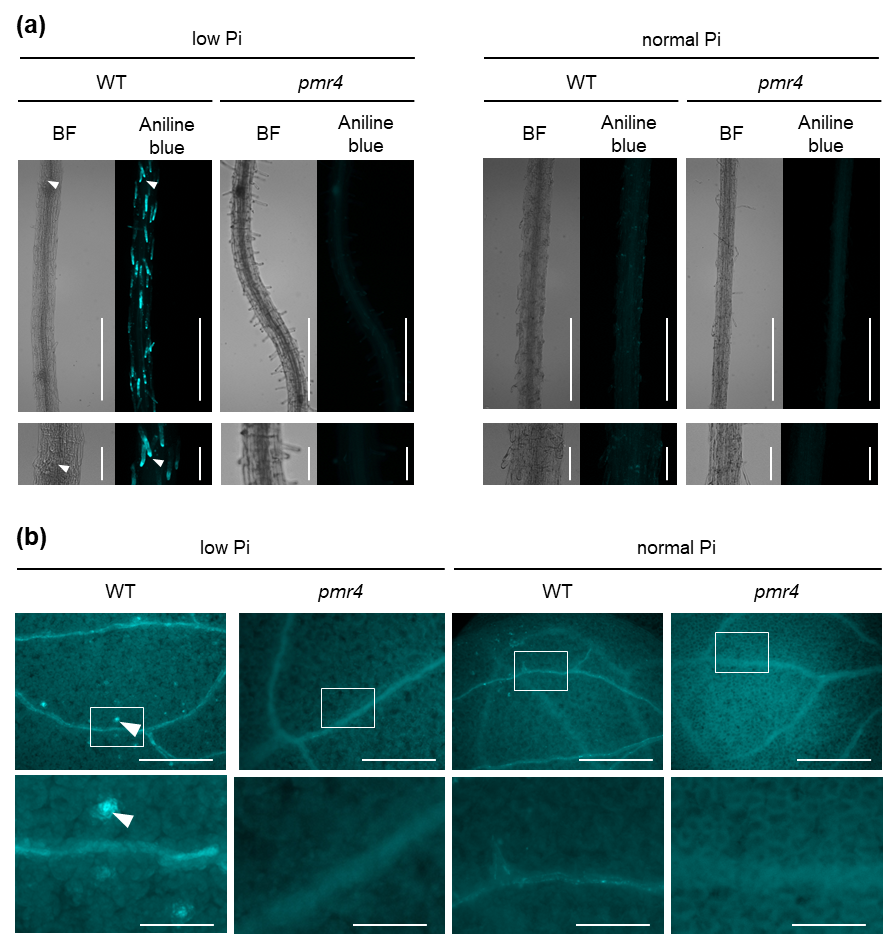

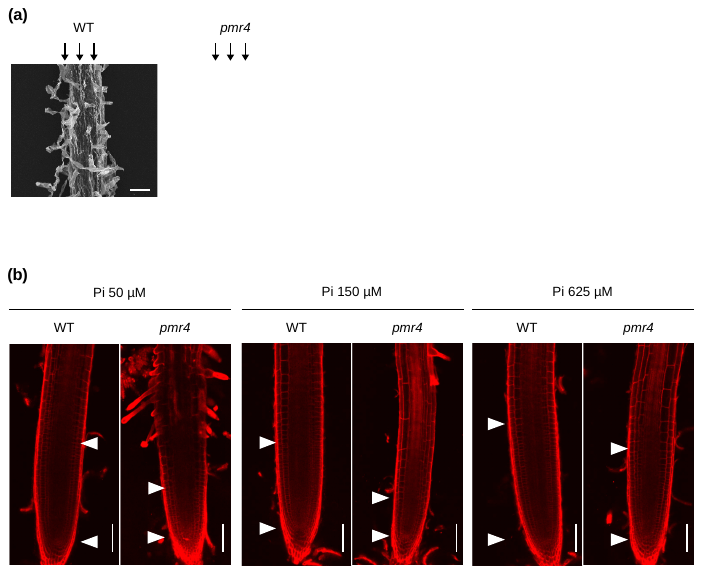

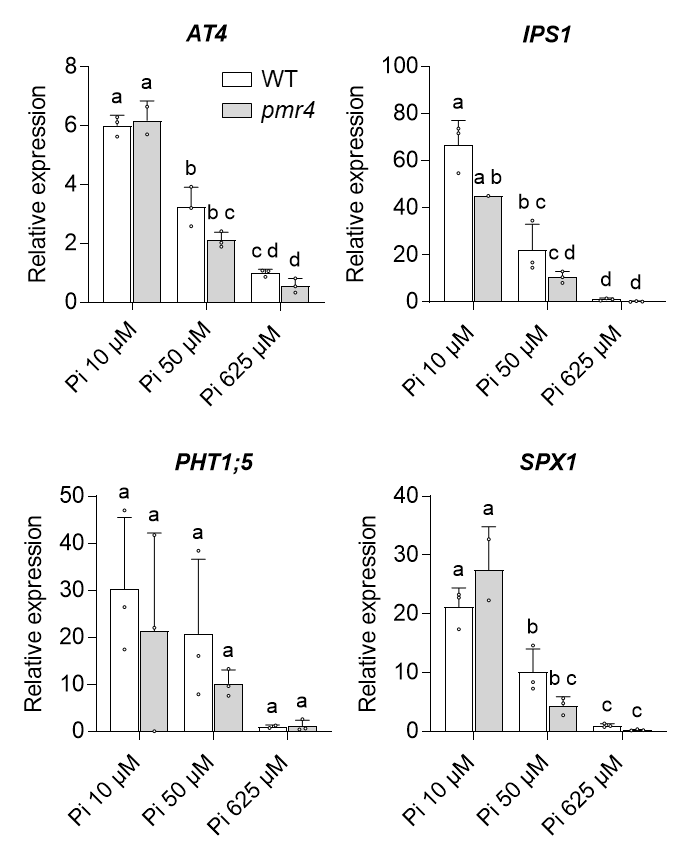

**Figure S3. Pi deficiency-induced callose deposition under dark conditions.**

Aniline blue staining of primary roots (a) and cotyledons (b) 3 d after transfer of 5-d-old *Arabidopsis thaliana* seedlings to low Pi (P 10 μM) or normal Pi (P 625 μＭ) under dark conditions. Magnified images of the square area in the upper panels are shown in the lower panels. White arrowheads indicate callose deposits. Bars, 500 µm (a, upper panel), 100 μm (a, lower panel), 500 µm (b, upper panel) and 100 μm (b, lower panel).

**Figure S4. Root hair and meristem morphology in *pmr4* under Pi deficiency.**

(a) Scanning electron microscopy images of the primary roots 10 d after transfer of 5-d-old *Arabidopsis thaliana* seedlings to low Pi (50 µM) liquid media. Arrow lines show vertical position of root hairs. Bars, 50 μm. (b) Meristematic region is reduced in *pmr4* roots under phosphate deficiency. Six-day-old *Arabidopsis thaliana* seedlings exposed to the indicated Pi media for 9 d were subjected to PI staining. Meristematic zone borders are indicated by white arrowheads. Bar, 100 μm**.**

**Figure S5. PSR marker gene expression in *pmr4* roots**

mRNA levels for PSR marker genes (*AT4, IPS1, PHT1;5* and *SPX1*) in roots of 7-d-old *Arabidopsis thaliana* plants, relative to the reference gene *ACT2,* determined by RT-qPCR analysis. Three-day-old seedlings were exposed to low Pi (10 µM or 50 µM) or normal Pi (625 µM) media for 4 d. Data show means ± SD (n = 2-3). Different letters indicate significant differences in two-way ANOVA and Tukey HSD test (*P* < 0.05).

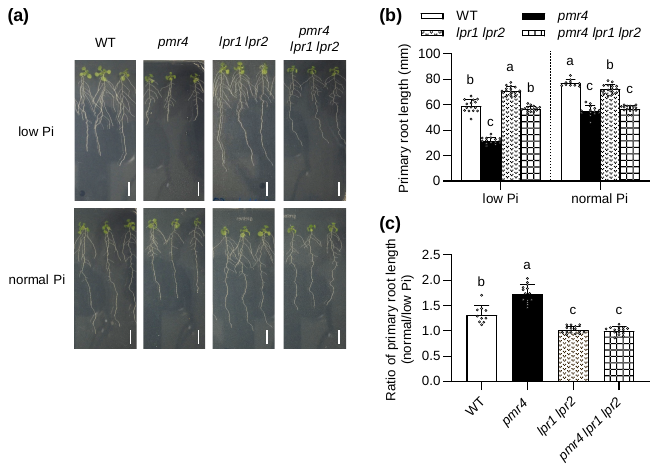

**Figure S6. *PMR4* restricts *LPR1/LPR2*-mediated inhibition of the primary root growth under low Pi.**

(a) Six-d-old *Arabidopsis thaliana* seedlings were exposed to low Pi (150 µM) or normal Pi (625 µM) media for 6 d. Bar, 10 mm. (b) Primary root length in (a). (c) Ratio of the primary root length in normal Pi relative to low Pi of (b). Data show means ± SD (n = 10~15). Different letters indicate significant differences in one-way ANOVA and Tukey HSD test (*P* < 0.05).

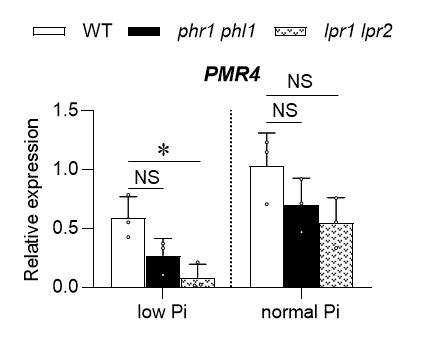

**Figure S7. *PMR4* expression in *phr1 phl1* and *lpr1 lpr2* roots.**

*PMR4* levels in roots of 7-d-old plants determined by RT-qPCR analysis relative to the reference gene *ACT2*. Three-day-old seedlings were exposed to low Pi (10 µM) or normal Pi (625 µM) media for 4 d. Data show means ± SD (n = 3). Asterisks (* *P* < 0.05) indicate statistically significant differences relative to WT according Dunnett’s multiple comparison test. NS, not significant.

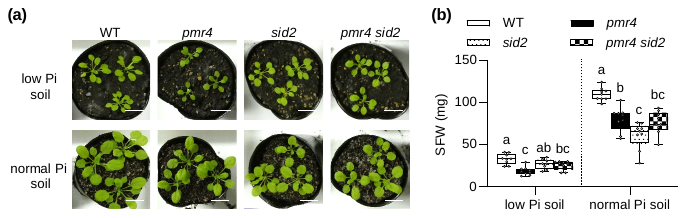

**Figure S8. Plant growth is lowered in *pmr4* on soil.**

(a) Ten-d-old *Arabidopsis thaliana* seedlings grown in low Pi and normal Pi soil for 14 d. Bars, 2 cm.

(b) Shoot FW in (a). Data show means ± SD (n = 8~9). Different letters indicate significant differences in one-way ANOVA and Tukey HSD test (*P* < 0.05).

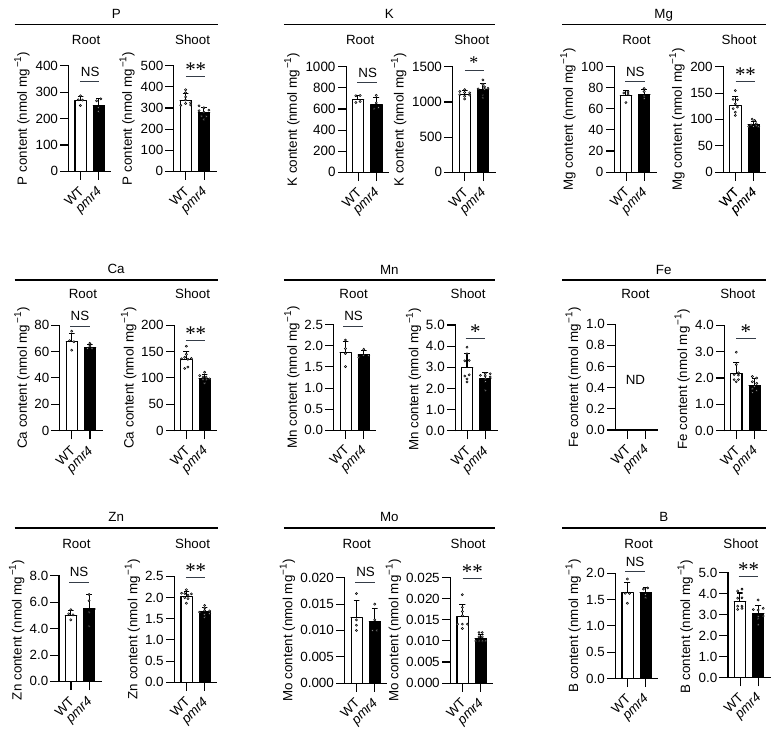

**Figure S9. Plant element content in shoot and root under normal phosphate conditions.**

Nutrient contents (P, K, Mg, Ca, Mn, Fe, Zn, Mo and B) per root or shoot dry weight of 5-d-old *Arabidopsis thaliana* seedlings exposed to normal Pi (625 µM) for 4 d. Data show means ± SD (n = 4~8). Asterisks (** *P* < 0.01, * *P* < 0.05) indicate statistically significant differences relative to WT according to Student’s *t*-test. NS, not significant. ND, not detected.

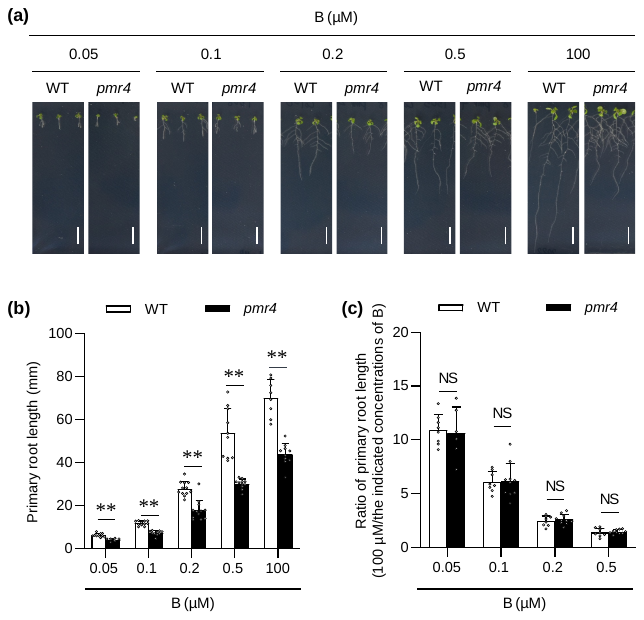

**Figure S10. Primary root growth inhibition under B deficiency.**

(a) Nine-d-old *Arabidopsis thaliana* seedlings were grown on the indicated concentrations of B. Bar, 10 mm. (b) Primary root length in (a). (c) Ratio of primary root length in the indicated B concentrations relative to B sufficiency at 100 µM of (b). Data show means ± SD (n = 6~12). Asterisks (** *P* < 0.01) indicate statistically significant differences relative to WT according to Student’s *t*-test. NS, not significant.

**Table S1. *Arabidopsis thaliana* mutants used in this study.**

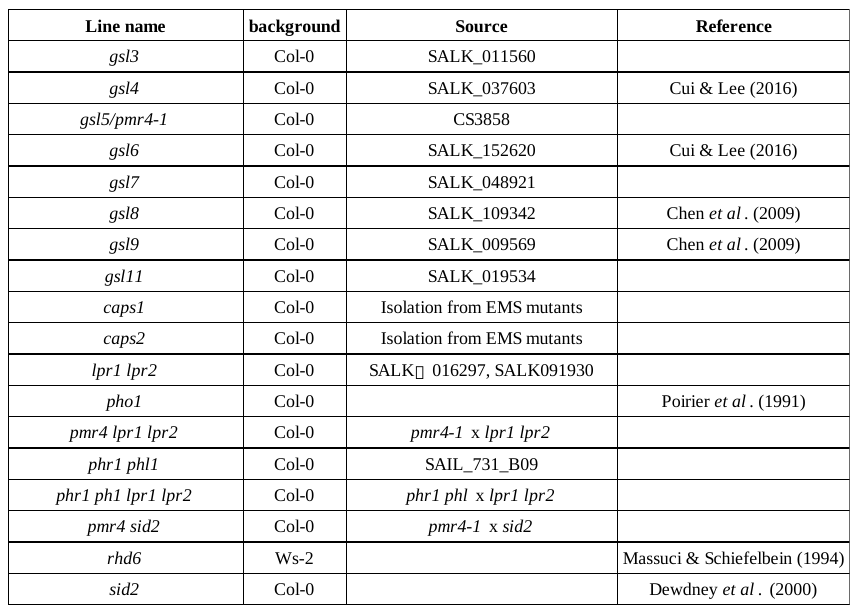

**Table S2. DNA primers used in construction and qPCR analyses**.
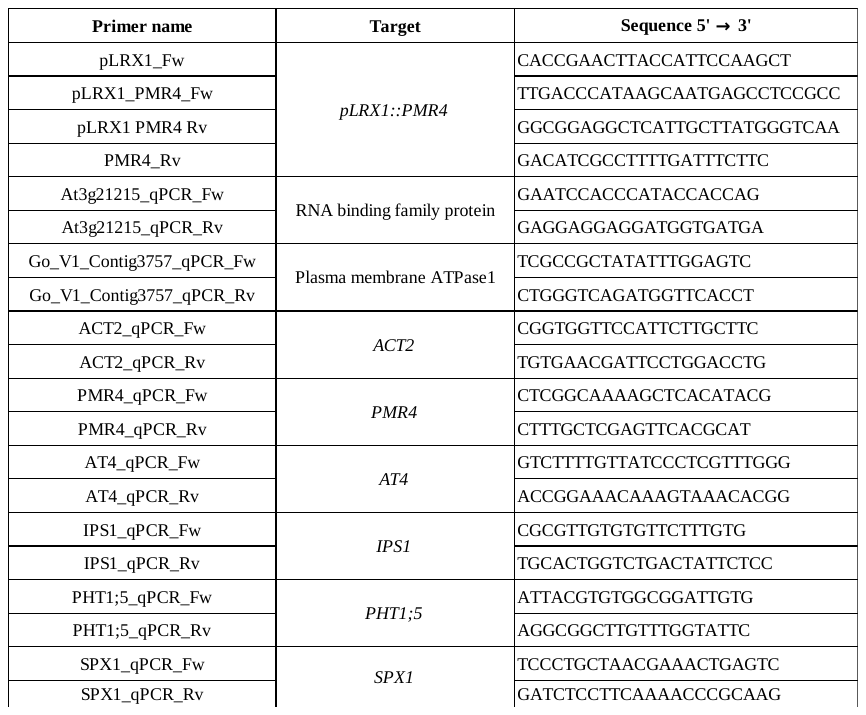
